# *In vivo* single-cell CRISPR screening for microproteins identifies a critical ribosomal component

**DOI:** 10.1101/2025.03.17.643322

**Authors:** Fabiola Valdivia-Francia, Umesh Ghoshdastider, Peter F. Renz, Daniel Spies, Mark Ormiston, Katie Hyams, Chenke Shi, Clara Duré, Merve Yigit, David Taborsky, Ameya Khandekar, Ramona Weber, Homare Yamahachi, Stephanie J. Ellis, Ataman Sendoel

## Abstract

The dark proteome includes a rapidly expanding catalog of microproteins with unknown functions that have been historically ignored in genome annotations. Here, we exploit an *in vivo* single-cell CRISPR screening strategy in the mouse epidermis to systematically investigate the tissue-wide function of microproteins. We document the global and cell-type-specific roles of microproteins during epidermal development and homeostasis at single-cell transcriptomic resolution. Focusing on select candidates, we identify a novel microprotein on *Gm10076*, identical to the ribosomal intersubunit bridge protein RPL41, whose perturbation strongly impairs proliferation and protein synthesis. Employing ribosome profiling and RNA sequencing, we show that *Gm10076* perturbation profoundly reshapes the translational landscape. Contrary to its prior classification as nonessential, we find that the ribosomal protein RPL41 is essential for cellular proliferation, warranting further investigation into its role as an intersubunit bridge in the translational machinery. Together, our study comprehensively charts the tissue-wide functional landscape of the dark proteome, uncovers a second *Rpl41* gene critical for ribosome function and establishes a basis for exploring the impact of microproteins on disease pathogenesis.

## Introduction

The mammalian genome encompasses numerous non-canonical microprotein-encoding open reading frames, which have been historically overlooked in genome annotations. Traditional annotation methods for protein-coding open reading frames (ORFs) in genomes focused on specific criteria, such as minimum length standards or amino acid conservation. This led to the exclusion of proteins shorter than 100 amino acids, which were dismissed as nonfunctional. However, recent genome-wide ribosome profiling and mass spectrometry studies have challenged the traditional annotation of ORFs, resulting in the identification of more than 7200 human noncanonical ORFs that encode microproteins, thereby substantially expanding the protein-coding capacity of a cell (*1, 2*).

Although the number of annotated microproteins is rapidly increasing, only a small fraction have been functionally characterized (*2, 3*). Microprotein function has been reported in diverse cellular processes, highlighting their biological activity in cellular signaling, gene regulation and metabolism, raising new questions about their significance as functional components of the mammalian genome (*4–8*). Additionally, microproteins may play widespread roles in driving the pathogenesis of diseases such as cancer, highlighted by pan-cancer analyses with thousands of microprotein transcripts deregulated in cancer (*4, 9*). However, the extent to which microproteins impact cellular function and how these roles vary in a cell-type-specific manner remains largely unknown. Understanding the contributions of microproteins to tissue function necessitates robust *in vivo* platforms that can systematically map microprotein functions in specific cell types.

Pooled CRISPR screens represent a powerful strategy to investigate the function of microproteins and have recently spurred the identification of functionally relevant microproteins (*10–14*). Although highly effective, pooled CRISPR screens have two major limitations. First, they are restricted to simple cellular readouts such as proliferation and cannot capture more complex cellular traits such as transcriptomics, which are among the most comprehensive characterizations of cellular function. Assessing transcriptomic changes may be particularly important for dissecting potentially more subtle functions of microproteins, which may remain undetectable through conventional proliferation-based assays. Second, pooled CRISPR screens are unable to assess cell-type-specific roles of microproteins. Uncovering cell-type-specific microprotein functions could profoundly enhance our understanding of how microproteins contribute to the complexity of mammalian proteomes.

Here, we exploit the mouse epidermis as a model to systematically dissect the tissue-wide function of microproteins at single-cell transcriptomic resolution. We employ ultrasound-guided *in utero* microinjections of lentiviral libraries into embryonic stage (E) 9.5 embryos and combine it with an adjusted CRISPR droplet sequencing (CROP-seq) strategy to document the cell-type-specific function of microproteins (*15, 16*). We unveil a second functional *Rpl41* gene encoded from a long noncoding RNA (lncRNA), which is – redundantly with the canonical *Rpl41 –* essential for cellular proliferation.

## Results

### Annotation of noncanonical ORFs from ribosome profiling data

To annotate translated noncanonical ORFs in healthy tissues and cancer, we first set out to analyze our own and published sets of ribosome profiling data of the mouse skin (*17*). We exploited the ORF regression algorithm for translational evaluation of ribosome-protected footprints (ORF-RATER), a ribosome profiling algorithm that uses a linear regression model to match ribosome occupancy patterns to expected profiles (*18*), thereby determining all possible ORFs with canonical and near-cognate start codon regardless of minimum length. Using the ORF-RATER algorithm, we annotated ORFs in both wild-type and pre-oncogenic SOX2-expressing mouse epidermis *in vivo*, as well as in wild-type keratinocytes and squamous cell carcinoma cells *in vitro* (Fig. 1A, Table S1). Furthermore, we included as an additional layer of evidence mapped translation initiation sites from harringtonine-treated samples, which blocks initiating but not elongating ribosomes (*19*) (Fig. 1A).

**Fig. 1:**
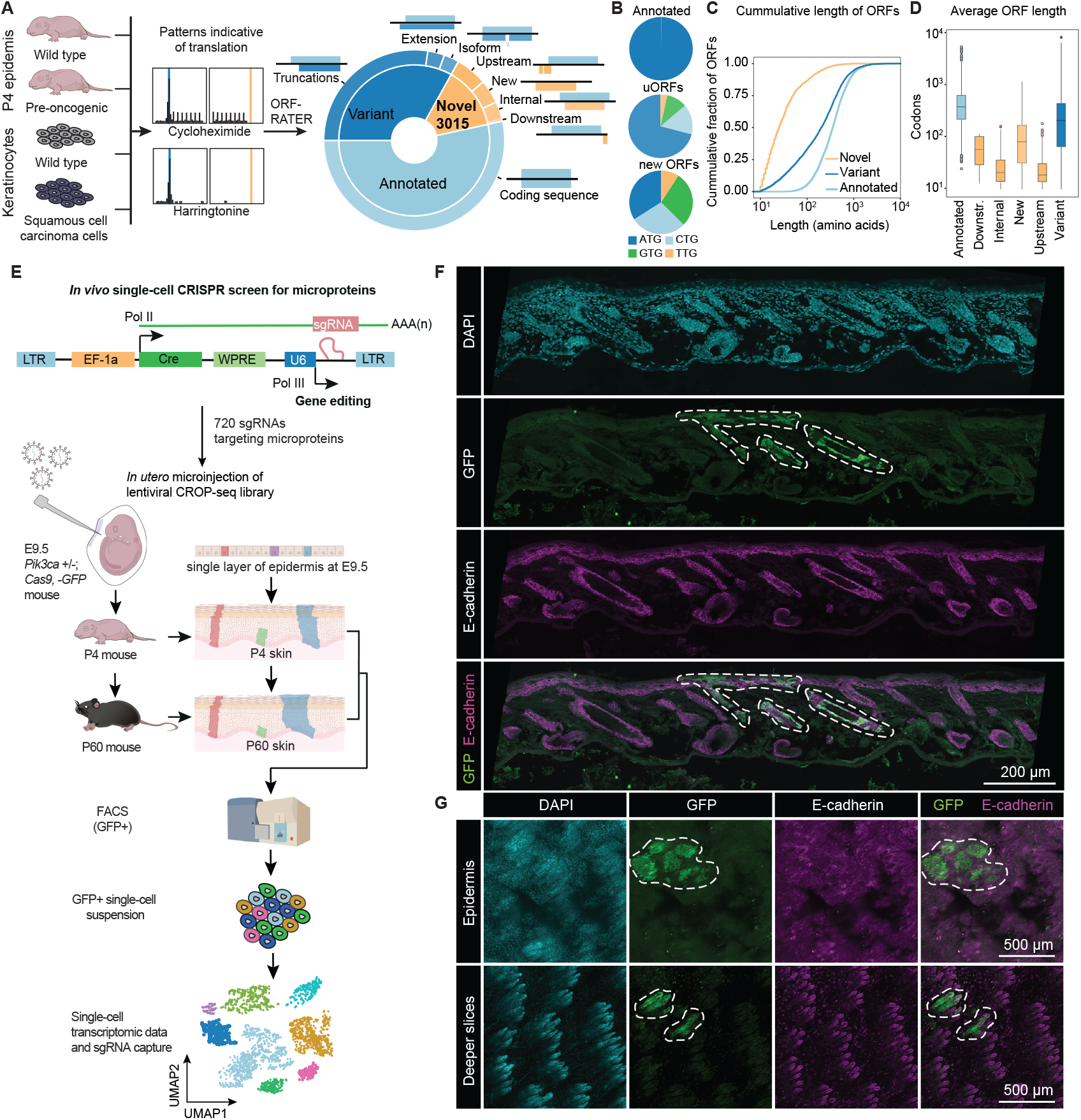
Annotation of microprotein-encoding noncanonical ORFs. **(A)** Workflow for annotating noncanonical ORFs from mouse ribosome profiling datasets. ORFs were classified using the ORF-RATER pipeline into three main categories: annotated ORFs, variant ORFs, and novel ORFs. **(B)** Pie charts showing the percentage of ORFs with ATG, CTG, GTG, or TTG start codons across different ORF categories. **(C)** Cumulative fraction plot showing the length distribution of ORFs. 76% of novel ORFs are shorter than 100 amino acids. **(D)** Box plot comparing the average lengths of annotated canonical ORFs (338 codons), variant ORFs (240 codons), novel ORFs (43 codons) and uORFs (15 codons). **(E)** Schematic of the single-cell CRISPR screen to identify functional microproteins. A pooled library containing 720 sgRNAs was injected in utero into the amniotic cavity of E9.5 *Pik3ca*^*H1047R+/-*^ *Cas9*^*+/-*^ embryos. Skin was harvested at postnatal day 4 (P4) and day 60 (P60), and GFP-positive cells were sorted for single-cell RNA sequencing and sgRNA capture. **(F)** Distribution of GFP-positive clones in sections of the P60 epidermis. E-cadherin staining marks epithelial cell junctions. Dashed lines mark GFP-positive clones. **(G)** Wholemount immunofluorescence of GFP-positive clones in the P60 mouse epidermis. E-cadherin staining marks epithelial cell junctions.

Applying ORF-RATER, we identified a total of 22,404 ORFs, among which 12,011 (54%) were recognized as annotated canonical open reading frames. 7378 (33%) of the identified ORFs were variants of annotated ORFs. Finally, we identified 3015 (13%) novel ORFs, including 1269 upstream ORFs (uORF), 851 new ORFs, 779 internal and 17 downstream ORFs (Fig. 1A). The representative ribosome profiling traces from newly identified microproteins showed similar footprint patterns as those from annotated coding sequences (CDSs), including the presence of triplet-nucleotide periodicity as detected by ORF-RATER, which is a hallmark feature of translation (Fig. S1A-D).

In the ORF-RATER annotation, 99.8% of canonical CDSs initiated with an AUG codon, while 49% of the novel ORFs utilized near-cognate start sites (NUG, where N represents C, G or U), with CUG being the most prevalent start codon (Fig. 1B). For uORFs, 29% start with a near cognate start codon, while 66% of the new ORFs initiate with non-AUG codons. Canonical ORFs had the longest median length at 338 codons, followed by variant ORFs with a median length of 240 codons and novel ORFs, which had a median length of 26 codons. Within novel ORFs, median lengths ranged from 15 codons for uORFs, 21 codons for internal ORFs, 43 codons for newly identified ORFs, up to 59 codons for downstream ORFs (Fig. 1C-D).

Prompted by the predicted translation of 3015 (13%) novel ORFs, we set out to investigate the function of the identified uORFs and new ORFs (herein both referred to as microproteins). We curated a high-confidence subset of predicted microproteins using the ORF-RATER rank sum approach, which ranks ORFs based on translation features via random forest classification (Table S1). Given their potential relevance to cancer, we prioritized microproteins highly translated in pre-oncogenic and oncogenic genetic backgrounds. This selection included a total of 156 uORF and 65 new ORFs. Additionally, we included six known uORFs within *Atf4, Mief1, Mdm2, Ctnnb1, Hras, and Pten*. Finally, 50 nontargeting single-guide RNAs (sgRNAs) were included as controls.

### *In vivo* single-cell CRISPR to target microproteins

To test the function of these microproteins, we sought to exploit our recently established *in vivo* single-cell CRISPR screening strategy to infer tissue-wide gene function in the mouse epidermis (*15*). This technique leverages ultrasound-guided microinjections of lentiviral libraries into E9.5 embryos and enables us to link the perturbation of microproteins in the mouse epidermis to its single-cell transcriptomic response (*15, 20, 21*) (Fig. 1E). The system capitalizes on the integration of a polymerase II-driven transcript encoding the sgRNA, facilitating direct guide capture during single-cell RNA sequencing (*16*). To explore the roles of annotated microproteins within a cancer-relevant genetic context, we employed an inducible *Pik3ca* mouse model. This model activates the H1047R hotspot mutation common in various human cancers, including head and neck and skin squamous cell carcinomas (*22, 23*) (Fig. 1E) and has been previously used as a sensitized oncogenic background for screens (*24, 25*). We then crossed this cancer model with the inducible Cas9-EGFP mouse to allow gene editing of microproteins. For recombination, we substituted the Puromycin cassette in the original CROP-seq vector with a Cre recombinase, thereby enabling simultaneous Cre-dependent recombination and expression of oncogenic PIK3CA and CAS9 in infected cells (Fig. 1E). Thus, our experimental setup allows the induction of an oncogenic background in combination with CRISPR-based gene editing, while infected cells can be tracked via green fluorescent protein (EGFP) expression (Figs. 1E-G, S1E).

To determine the function of microproteins, we subcloned the library of 720 sgRNAs targeting 229 microproteins into our adjusted CROP-seq Cre vector, with three sgRNAs per microprotein (Fig. 1E, Table S2). However, due to the scarcity of NGG motifs required for the CRISPR system, some microproteins shorter than 45 nucleotides were targeted with only two sgRNAs (Table S2). Given the limited design options for selecting high-efficiency sgRNAs, we tested the editing efficiency of 45 sgRNAs. While 21 out of 45 demonstrated robust editing (>25% editing efficiency), their overall efficiency was, as expected, lower than that typically observed for sgRNAs targeting conventional genes (*15*) (Fig. S2). We then produced high-titer lentivirus and used the *in utero* ultrasound-guided microinjection system to deliver the library into the amniotic cavity of E9.5 mouse embryos. At this stage in development, there is only a single layer of epidermis, which allows for the lentivirus to infect the ectoderm surface progenitors and is consequently stably propagated during development. To target each cell with a single sgRNA-containing virus particle, we chose infection rates between 2-10% (Fig. 1F-G), which are below the expected multiple infection range and minimize the likelihood of double-infected cells (*20, 21*). We collected the mouse epidermis at postnatal day 4 (P4) or day 60 (P60) and isolated EGFP-positive, sgRNA-containing cells using fluorescent-activated cell sorting (FACS). These cells were then processed to generate single-cell RNA sequencing (scRNA-seq) libraries alongside sgRNA capture (Fig. 1E).

We generated scRNA-seq libraries of a total of 700,726 cells across the P4 and P60 time points. Following stringent filtering for cells with low unique molecular identifier (UMI) counts and for cells without sgRNA capture, we obtained 196,704 P4 epidermal cells and 197,581 P60 cells expressing a single sgRNA. This accounts for a robust coverage of 273 (P4) and 344 (P60) cells on average per sgRNA (Fig. 2A-G, S3). As previously shown (*15*), the *in vivo* single-cell CRISPR screening approach enables us to detect the major epidermal cell populations, including *Krt14*-positive epidermal stem cells, *Krt10*-positive suprabasal cells, upper hair follicles, inner and outer bulge stem cells, sebaceous glands and immune cells such as T cells, macrophages and dendritic cells (Fig. 2B-G).

**Fig. 2:**
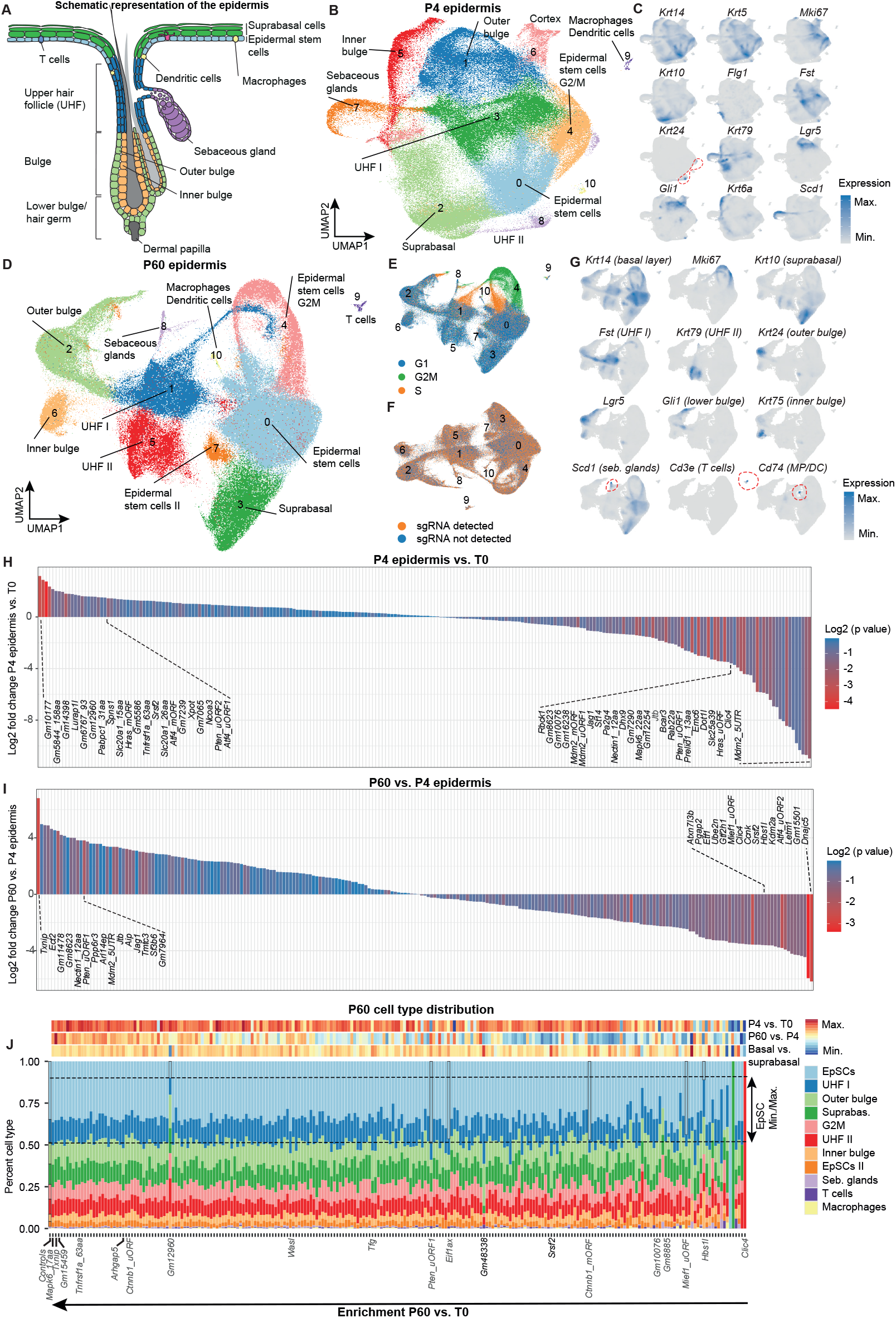
*In vivo* single-cell CRISPR screen for microproteins. **(A)** Schematic representation of the different cell populations in the P60 mouse epidermis. **(B)** UMAP of the 11 cell populations identified in the P4 epidermis (n= 196,704 cells with sgRNA capture from 18 mice). UHF, upper hair follicle. **(C)** Cell-type specific markers gene expression by Nebulosa density plots for the P4 epidermis. **(D)** UMAP of the 11 cell populations identified in the epidermis of P60 mice (n= 197,581 cells from mice after filtering for annotated sgRNAs). UHF, upper hair follicle. **(E)** Cell cycle phase distribution across cell types at P60. **(F)** sgRNAs are uniformly detected across cell types. UMAP depicting total GFP-sorted cells at P60, including cells with and without sgRNA annotation. **(G)** Nebulosa density plots of cell type-specific marker gene expression in the P60 epidermis. Red circles highlight Scd1-positive sebaceous gland cells, Cd3e-positive T cells, and Cd74-positive macrophages/dendritic cells (MP/DC). UHF, upper hair follicle. **(H)** Waterfall plot showing the log2 fold change in sgRNA representation in the P4 epidermis compared to the library (T0), computed with MAGeCK and normalized to 50 nontargeting sgRNAs. Bars are colored by p value, highlighting the top enriched and depleted microprotein candidates. **(I)** Waterfall plot showing the log2 fold change in sgRNA representation in the P60 epidermis compared to P4. **(J)** Cell type distribution of the 229 perturbed microproteins in the P60 epidermis, sorted by enrichment at P60 compared to library (T0). Maximum and minimum percentage of epidermal stem cells (EpSCs) is depicted by dashed lines. Dashed lines indicate maximum and minimum percentages of epidermal stem cells (EpSCs). The black box highlights microproteins with distinct differences in EpSC proportions. Heatmaps display sgRNA representation changes for P4 vs. T0, P60 vs. P4, and basal vs. suprabasal ratios for the 229 microproteins.

The epidermis provides a powerful model for studying tissue dynamics and the intricate balance between proliferation and differentiation (*21, 26, 27*). At P4 stage, the strategy enables the identification of microproteins linked to embryonic development. From P4 to P60, following the completion of epidermal development, we can explore microproteins affecting epidermal tissue homeostasis, thereby bridging key developmental and homeostatic phases of epithelial biology. First, we assessed the overall effect of microproteins on epidermal proliferation using the conventional CRISPR screening readout, considering sgRNAs representation at P4 compared to the initial library (T0). While microprotein perturbations result in a loss-of-function of the respective microprotein, phenotypes observed upon uORF targeting can arise from two distinct mechanisms. They may result either from the loss of the uORF-encoded microprotein itself or from disruptions in uORF-mediated translational control of the main ORF. Among the top depleted P4 candidates, we found sgRNAs targeting the uORFs/5’UTR of several cancer-associated genes, including *Mdm2, Hras*, and *Pten* (Fig. 2H, S4). These cancer genes are known to harbor complex 5’UTRs. For example, the oncogene *Mdm2* is reported to toggle between two transcript isoforms, where one contains two uORFs, resulting in a poorly translated mRNA (*28*). Furthermore, the tumor suppressor *Pten* contains several N-terminal extensions and uORFs that also control the ratio of these proteoforms (*29–31*) (Fig. S1C). In contrast, perturbation of the microproteins *Gm10177, Gm5844* and *Gm14398* led to a strong enrichment, suggesting that these microproteins negatively impact epidermal development. In addition, perturbation of *Gm7239*, a recently identified nuclear microprotein in the liver discovered through a mass spectrometry-based approach (*32*), was similarly among the top enriched micro-proteins (Fig. 2H).

Between P4 and P60 stages, after the stratification events during epidermal development, perturbations of the uORFs *Dnajc5, Letm1, Atf4* and *Gm15501* microprotein were markedly depleted (Figs. 2I, S4). Conversely, perturbations of the uORFs on *Txnip, Ect2* and microproteins on *Gm11478* and *Gm8623* were strongly enriched between the P4 and P60 stages. The thioredoxin-interacting protein *Txnip*, known to harbor a previously described uORF (*33*), is a multifunctional protein implicated in oxidative stress response, stress-induced apoptosis and pyroptosis (*34, 35*). On the other hand, epithelial cell transforming sequence 2 (*Ect2*) is a guanine nucleotide exchange factor (GEF) for the Rho GTPases *Rac1, Cdc42* and *RhoA* and is frequently overexpressed in cancer (*36*), suggesting that loss of uORF-mediated translational control of *Ect2* may result in epithelial hyperproliferation (Fig. 2I).

Exploring the cell type distributions of the microprotein perturbations at P60, we found that uORF targeting of *Pten, Mief1* and *Eif1ax* led to the highest relative expansion of epidermal stem cell populations, indicating that these uORFs may specifically impact epidermal stem cell proliferation (Fig. 2J). Furthermore, zooming into epidermal differentiation and assessing sgRNA ratios in epidermal stem cells compared to suprabasal cells, we observed that the perturbations of *Gm48338* and of the uORFs of *Tfg, Srsf2* and *Wasl* exhibited among the highest anti-differentiation index (Fig. 2J, sgRNAs epidermal stem cells/suprabasal). The actin cytoskeleton-remodeling protein WASL (N-WASP) is essential for epidermal differentiation (*37*) and skin tumor formation in a p53-dependent manner (*38*), suggesting that perturbation of the *Wasl* uORF and potential uORF-dependent alteration of WASL protein levels may underlie the differentiation block of epidermal stem cells.

Collectively, our single-cell microprotein CRISPR screening across the developmental and homeostatic phases of the epidermis suggests that a surprisingly large cohort of microproteins annotated from our ribosome profiling data play cellular roles *in vivo* and provide a curated list of microproteins for detailed mechanistic exploration.

### Charting microprotein function at single-cell transcriptomic resolution

The transcriptome provides the most comprehensive characterization of cellular states and may unveil microprotein functions that are not captured by classic pooled CRISPR screens. Given that the core strength of our *in vivo* single-cell CRISPR screening strategy lies in its transcriptomic data, we next explored the differentially expressed genes upon perturbation of the 229 microproteins in each of the 11 P60 cell types (Table S3). Remarkably, a cohort of 22 microprotein perturbations triggered substantial transcriptional changes, with each inducing over 500 differentially expressed genes in epidermal stem cells (Fig. 3A, marked in red). These findings suggest that this subset of microproteins profoundly shapes gene expression and cellular function.

**Fig. 3:**
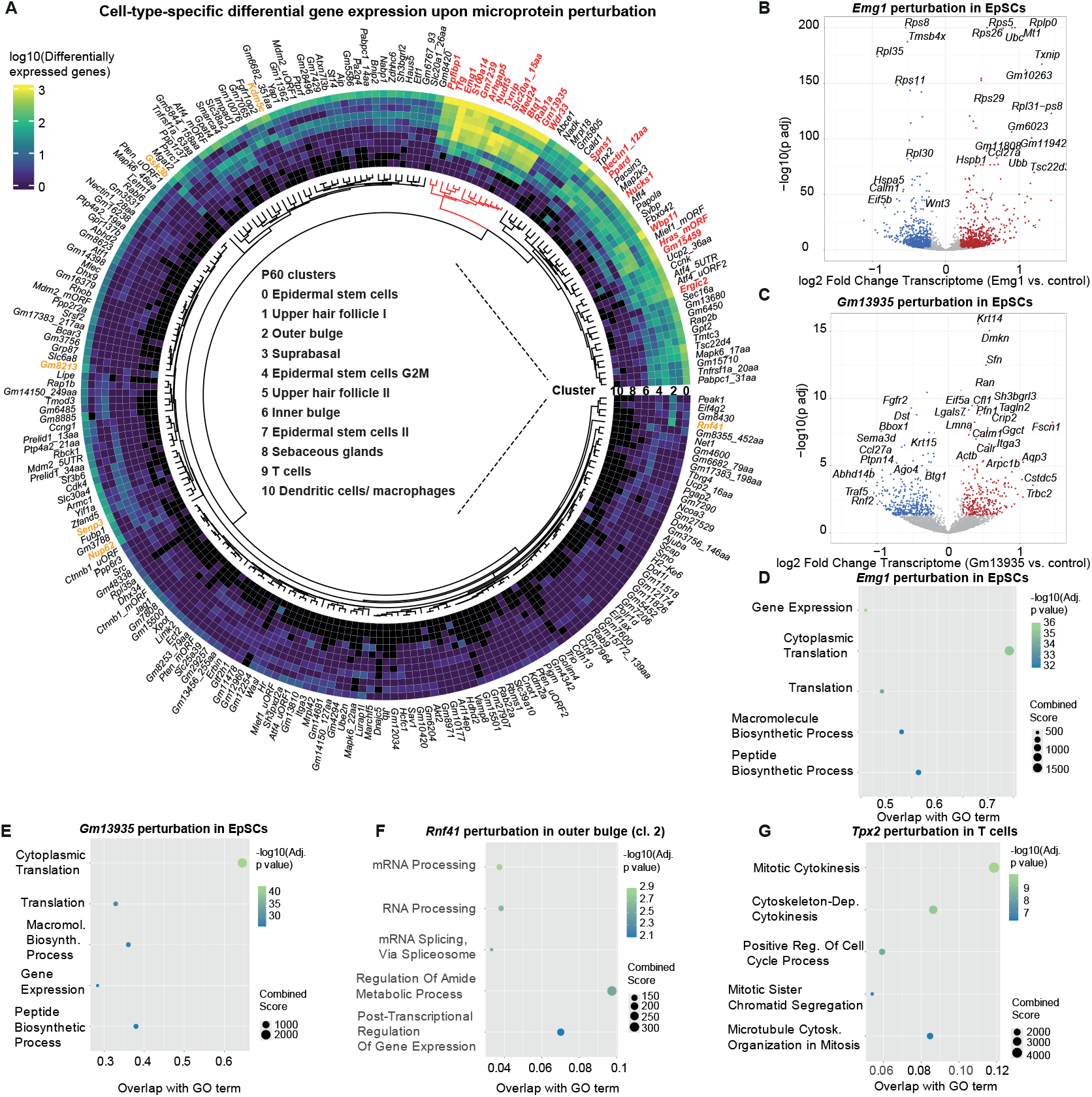
The translational landscape of microprotein perturbation. **(A)** Circos plot showing the number of differentially expressed genes for each of the 229 perturbations across the 11 cell populations within the P60 epidermis. Black sections indicate populations with insufficient cell numbers to determine differentially expressed genes. Perturbations that led to > 500 differentially expressed genes in epidermal stem cells are marked in red. Additional genes that resulted cell-type-specific differential expression are marked in orange. **(B)** Volcano plot displaying the top differentially expressed genes in epidermal stem cells upon perturbation of the *Gm13935* microprotein. **(C)** Volcano plot highlighting the top differentially expressed genes in epidermal stem cells upon *Emg1* uORF microprotein perturbation. **(D-E)** GO term enrichment analysis of differentially expressed genes in epidermal stem cells upon perturbation of the *Gm13935* and *Emg1* uORF microproteins. **(F)** GO enrichment analysis of DEGs in T cells upon *Tnfrsf1a_63* uORF perturbation, revealing activation of FAS signaling.

Generally, we observed both global and cell-type-specific alterations in transcript levels upon microprotein perturbation. A cluster of 14 microprotein perturbations induced substantial gene expression changes across at least 7 cell types (Fig. 3A, dendrogram marked in red). This cohort included *Arhgap5, Emg1, Txnip, Nudt5* uORFs as well as *Gm13935* and *Gm7239*, a nuclear microprotein previously described in the liver (*32*). Notably, disruption of the *Emg1* uORF led to widespread transcriptional changes, with 2809 differentially expressed genes in epidermal stem cells and 1556 in suprabasal (Fig. 3A-B). *Emg1* is required for pseudouridine methylation of the 18S rRNA and the biogenesis of the 40S ribosomal subunit (*39, 40*). Consistent with this function, ribosome, ribosomal biogenesis and cytoplasmic translation were the main enriched Gene Ontology (GO) terms in the differentially expressed genes (Fig. 3B, D). Similarly, perturbation of the 31 amino acid microprotein *Gm13935* led to 1344 differentially expressed genes in epidermal stem cells (Fig. 3C). *Gm13935* is a pseudogene of the ribosomal protein *Rps3a1*, almost fully aligning to amino acids 105 to 137 of *Rps3a1* (Fig. S1D). Perturbation of *Gm13935* similarly resulted in a strong enrichment of the GO term ribosome, suggesting a potential role for this truncated RPS3A1 microprotein in modulating ribosome function (Fig. 3E).

To further examine the impact of this cohort of 14 micro-protein perturbations, we performed an unbiased clustering analysis to group perturbations based on transcriptional similarity in P60 epidermal stem cells. This perturbation-perturbation matrix identified several distinct clusters, including a major cluster containing the ribosome biogenesis factor *Emg1*, alongside the nuclear microprotein *Gm7239, Tfg* and *Arhgap5* (Fig. S5A). Notably, all four perturbations exhibited a strong enrichment for downregulated ribosome biogenesis genes (Fig. S5B). For example, *Gm7239* perturbation led to the profound suppression of key ribosome assembly factors, including *Bms1* and *Utp* complex members for 40S biogenesis, *Rps2* and *Rrs1* involved in the 5S RNP complex and the snoRNP components *Nop56, Nop58* and *Dkc1* essential for rRNA modifications. This signature of ribosome biogenesis suppression upon *Gm7239* perturbation was conserved during differentiation, as indicated by the persistent enrichment of ribosome biogenesis in downregulated genes in suprabasal cells (Fig. S5C). The microprotein encoded on *Gm7239* is a pseudogene of SET (SET nuclear proto-oncogene), which, among other functions, acts as a histone acetylase inhibitor (*41*). Together, these findings indicate that this cohort of microproteins may share a common role in directly or indirectly modulating ribosome biogenesis.

We also observed cell-type-specific transcriptional alterations upon microprotein perturbation. Perturbation of *Senp3 Gm8213* and the lysine demethylase *Kdm5c* showed robust transcriptomic changes specifically in epidermal stem cells (Fig. 3A). Conversely, the E3 ubiquitin ligase *Rnf41* uORF led to gene expression alteration predominantly in the outer bulge of the hair follicle but not the other epidermal cell types (Fig. 3A). GO term analysis revealed strong enrichment in mRNA processing pathways, suggesting a potential outer bulge-specific role of *Rnf41* uORF in modulating mRNA splicing. This was evidenced by altered expression of several splicing factors, including *Sfpg, Sf3b3, Snrpd1* and *Snrpd3* (Fig. 3F). Furthermore, disruption of *Tpx2* uORF induced robust gene expression changes in T cells, with a notable enrichment of mitotic cytokinesis genes (Fig. 3A, G). TPX2 is a well-characterized microtubule-associated protein essential for mitotic spindle assembly (*42, 43*), highlighting the potential regulatory role of *Tpx2* uORF in modulating TPX2-dependent cytokinesis predominantly in T cells.

In summary, our cell-type-specific transcriptomic analyses chart the landscape of microprotein function and indicate that a small cohort of microproteins have profound and tissue-wide consequences on the cellular transcriptome.

### Validation screens for candidate microproteins

To validate the results of our *in vivo* single-cell CRISPR screen, we selected a sub-subset of microproteins to confirm their role in the epidermis (Fig. 4A-B). We chose 29 microproteins from different categories, including proliferative changes at P4, P60 or P60 versus P4, cell-type-specific effects and strong transcript level changes (Fig. S6). Additionally, we included 12 nontargeting sgRNA controls as well as 3 sgRNAs targeting *Rpl11* and *Tpr53* as proliferation controls (*15*). The total set of 101 sgRNAs was cloned into the same modified CROP-seq-Cre vector.

**Fig. 4:**
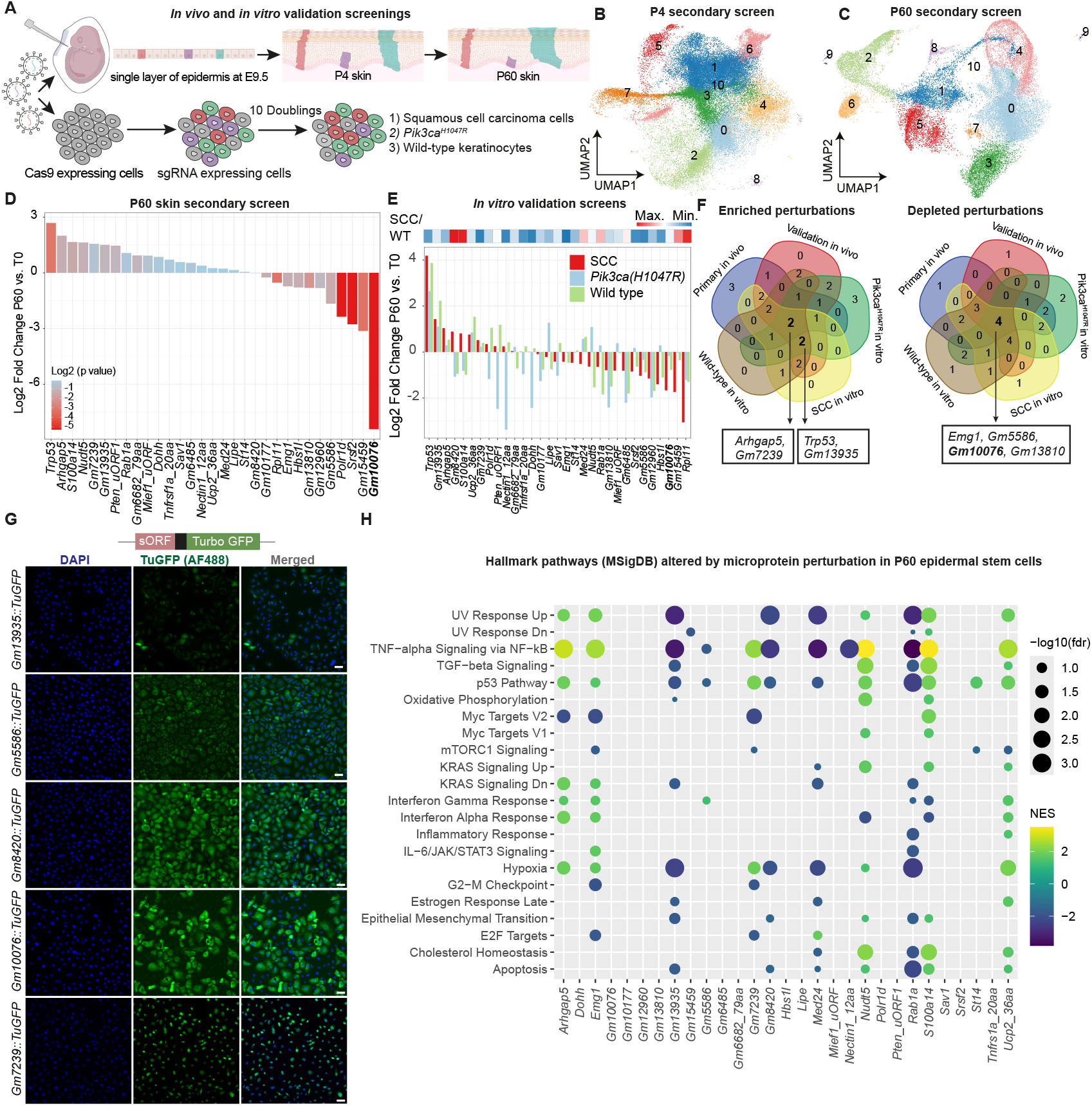
Validation screen identifies the microprotein *Gm10076*. **(A)** Schematic outline of the *in vivo* and *in vitro* CRISPR validation screens. For in vivo screen, pooled library containing 101 sgRNAs were injected *in utero* into *Pik3ca*^*H1047R+/-*^; *Cas9+/-* embryos at stage E9.5. Skin was collected at postnatal day 4 (P4) and 60 (P60) and sorted for GFP-positive cells for single-cell sequencing. For the *in vitro* screen, Cas9-expressing cells were infected with the library and passaged for 10 generations. **(B)** UMAP of the 11 cell populations identified in the skin of P4 mice (n= 58,307 cells from 3 mice, filtered for annotated sgRNAs). UMAP projections were generated together with the primary screen. **(C)** UMAP of the 11 cell populations identified in the skin of P60 mice (n= 32,390 cells from 2 mice, filtered for sgRNAs). UMAP projections were generated together with the primary screen. **(D)** Waterfall plot showing the log2 fold change in sgRNA representation in the P4 epidermis compared to the library (T0), computed with MAGeCK and normalized to 50 nontargeting sgRNAs. Bars are colored by p value. *Gm10076* perturbation showed the strongest depletion at P60. **(E)** Waterfall plot for the *in vitro* validation screen, highlighting the enriched and depleted microproteins in squamous cell carcinoma cells (SCCs), *Pik3ca*^*H1047R+/-*^ keratinocytes and wild-type keratinocytes, ordered by enrichment in SCCs. The heatmap shows the differences in sgRNA representation between SCC and wild-type keratinocytes. **(F)** Venn diagram showing the overlap of enriched and depleted microproteins across screens. *Arhgap5* and *Gm7239* are consistently enriched, while *Emg1, Gm5586, Gm10076* and *Gm13810* show consistent depletion across primary and validation screens. **(G)** Immunofluorescence and analysis of subcellular localization for 5 GFP-tagged microproteins. Scale bars, 50 μm. **(H)** Hallmark pathway activation (Molecular Signatures Database, MSigDB) following perturbation of the 29 candidate microproteins included in the validation screen, revealing key cellular processes regulated by the candidate microprotein. Gene Set Enrichment Analysis (GSEA) was performed for each of the 29 candidate microproteins in epidermal stem cells of the P60 epidermis. NES, normalized enrichment score in GSEA analysis.

First, we validated candidate microproteins *in vivo*. We collected 58,307 cells for the P4 timepoint (coverage of 550 cells per sgRNA) and 32,390 cells for the P60 timepoint (305 cells per sgRNA) (Fig. 4B-C). As expected, *Rpl11* perturbation led to depletion, while sgRNAs targeting *Trp53* resulted in a proliferative advantage (Fig. 4D). In keeping with the primary screen, perturbation of the uORFs of *Nudt5, Arhgap5, S100a14* and the truncated *Rps3a1*-like microprotein *Gm13935* showed a proliferative advantage. Among the depleted microproteins, we could confirm that perturbations of *Srsf2, Gm5586, Gm13810* and *Gm10076* exhibited a strong depletion, which is in line with the primary screening results. We also evaluated the differentially expressed genes upon microprotein perturbations in the secondary screen. For example, perturbation of *Gm13935* was among the microproteins with the strongest transcript level alterations (Fig. 3A). *Gm13935-*targeted cells also showed a high number of differentially expressed genes (669 genes in epidermal stem cells) in the validation screen, confirming the transcriptional data underlying the primary screening.

Second, we sought to also validate our microprotein candidates in an *in vitro* screen, as a confirmed role in keratinocytes would allow us to further examine microprotein function using *in vitro* assays. Additionally, the *in vitro* screen enabled us to test different genetic backgrounds. We selected three different cell lines and stably expressed CAS9: squamous cell carcinoma (SCC) cells (*17*), oncogenic *Pik3ca(H1047R*) keratinocytes established from our mouse model and wild-type keratinocytes (Fig. 4A, S7A). We then infected these 3 cell lines with the sgRNA library and cultured them for 10 passages, followed by sequencing of amplified sgRNAs. *Trp53* sgRNAs were the top enriched perturbations and *Rpl11* sgRNAs were among the most depleted, confirming the overall screening strategy (Fig. 4E). Examining sgRNA representation, we identified microproteins with potential cancer-specific functions, such as *Gm8420* or *S100a14* specifically in SCCs (Fig. 4E). In contrast, other microproteins such *Gm13935* and *Gm12960* showed consistent proliferative alterations across the 3 genetic backgrounds (Fig. 4E). To narrow down our candidates, we then compiled a high-confidence list of validated microproteins that consistently showed the same phenotype across the primary, secondary and *in vitro* screenings. These microproteins included the enriched perturbations of *Arhgap5, Gm7239* and the depleted perturbations of *Emg1, Gm5586, Gm13810* and *Gm10076* (Figs. 4F, S7B).

To test the translatability of our annotated microproteins, we next selected 5 microproteins and stably expressed them with a turbo GFP tag (Fig. 4G, S8). All 5 microproteins *Gm13935, Gm7239, Gm5586, Gm8420* and *Gm10076* were translated, albeit with different efficiencies, as also confirmed by western blots (Fig. S8A). Interestingly, these microproteins exhibited distinct subcellular localizations, including nuclear localization of *Gm7239* (Fig. S8B), consistent with its previous identification in liver nuclei (*32*), indicating that these microproteins are sufficient to alter the subcellular distribution of large fluorescent tags such as GFP. The expression level of *Gm13935*, whose sgRNAs were strongly enriched in our screen, was particularly low in expression in immunofluorescence and western blot analyses, even after sequential FACS-purification of GFP-high cells. This low GFP expression was dependent on *Gm13935* as a 2A cleavage peptide between the microprotein and GFP resulted in high GFP expression level across all cells (Fig. S8B). Given the high stability of turbo GFP, we surmised that the 31 amino acid microprotein must be actively degraded. Indeed, proteasomal inhibition by MG132 markedly increased turbo GFP expression in a time course (Fig. S8C-D). Together, our findings suggest that the annotated microproteins are efficiently translated and underscore their potential involvement in a broad spectrum of subcellular processes

Finally, to investigate in detail how the 29 candidate microproteins influence key cellular processes and potentially contribute to disease pathogenesis, including cancer, we performed hallmark pathway enrichment analysis on the differentially expressed genes (Fig. 4H). Several microproteins, including the Ras oncogene family gene *Rab1a, Emg1, S100a14 and Gm13935* exhibited widespread hallmark pathway changes, including p53 signaling, TGFβ, TNF or hypoxia signaling (Fig. 4H). Of note, several highly depleted microprotein perturbations, such as *Gm10076*, lacked enrichment in hallmark pathways (Fig. 4H). This raises the possibility that at P60, the remaining surviving cells are largely non-edited, resulting in minimal detectable transcriptomic changes. Collectively, these findings highlight the broad regulatory role of microproteins and underscore their impact on diverse cellular processes.

### *Gm10076* encompasses a second *Rpl41* gene

*Gm10076* perturbations surfaced as the top depleted candidate and were consistently depleted throughout our primary and validation screens (Fig. 4D-F). The 25 amino acid microprotein encoded on chromosome 14 matches the exact RPL41 protein sequence, indicating the potential existence of a second *Rpl41* gene in the mouse genome (Fig. 5A-B). Despite its name, *Rpl41* is neither associated with the 40S or 60S subunits, but constitutes a bridge protein positioned at the intersubunit interface of the 80S ribosome close to the decoding center, with contacts to rRNA elements of both ribosomal subunits (*44*) (Fig. 5C). Interestingly, based on studies in yeast and cancer dependency maps (*45, 46*), *Rpl41* has been regarded as a nonessential ribosomal protein associated with minimal phenotypic effects. Given the consistent and strong phenotype of *Gm10076* perturbation, which may surpass that of *Rpl41* mutations (*45, 46*), we sought to further investigate its functional involvement in the mammalian ribosome.

**Fig. 5:**
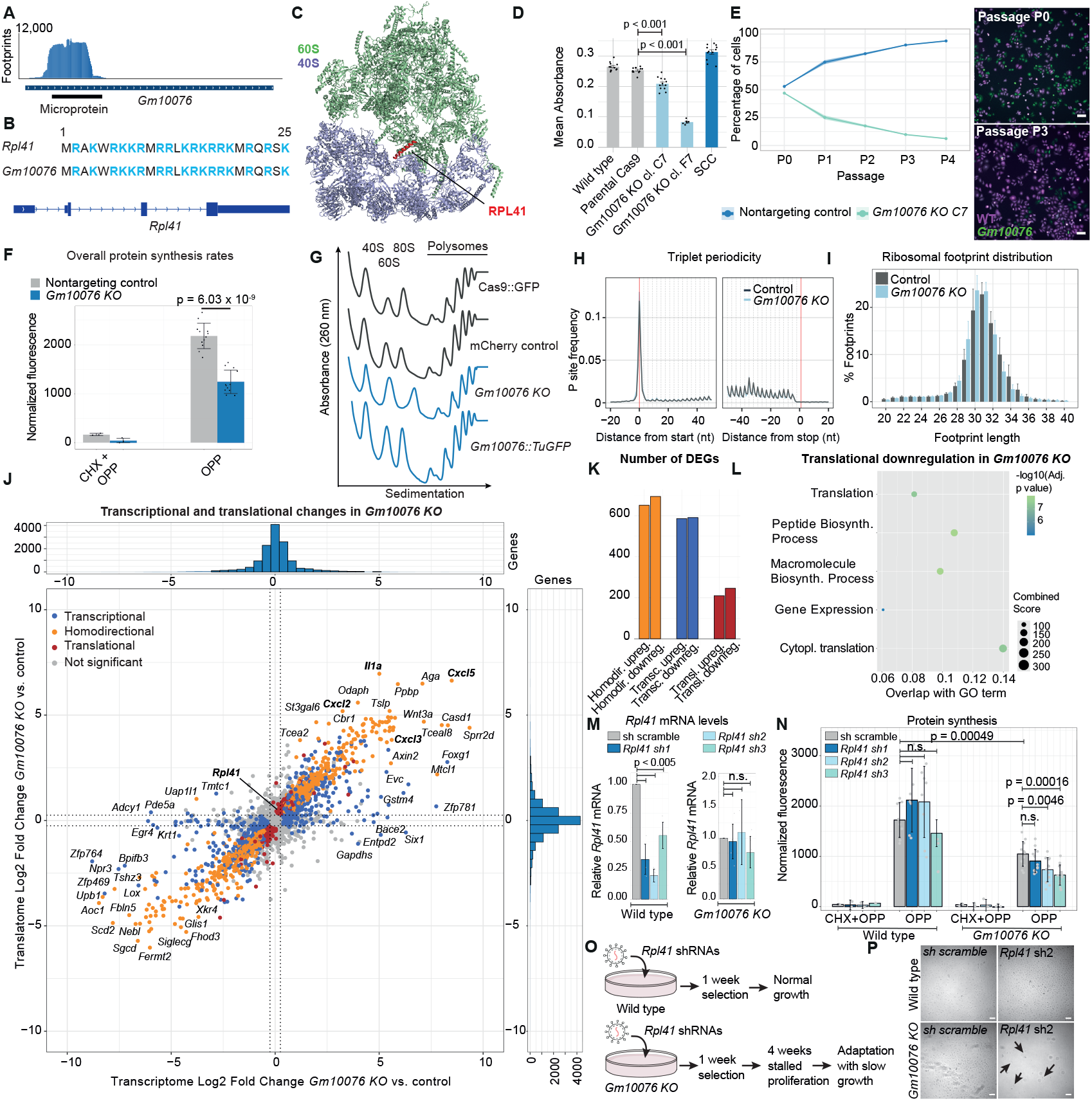
*Gm10076* encodes a second RPL41 gene and strongly shapes the translational landscape. **(A)** Ribosome profiling identifies the translation of the 25-amino acid microprotein on the lncRNA Gm10076 on chromosome 14. **(B)** Amino acid alignment of RPL41 and Gm10076 microprotein, highlighting positively charged residues in blue, which constitute 68% of the sequence. The lower panel illustrates the genomic structure of the canonical *Rpl41* gene, including introns and exons. **(C)** Structure of the mammalian cytosolic 80S ribosome (Oryctolagus cuniculus) highlighting RPL41 (red) at the intersubunit bridge. The 60S subunit is shown in green, the 40S subunit in light blue. **(D)** Proliferation analysis using an MTT assay comparing wild-type, parental *Cas9::GFP, Gm10076 KO C7, Gm10076 KO F7* and squamous cell carcinoma (SCC) cells. Data from 12 replicates (6 replicates for the F7 clone) are shown, with error bars representing the standard deviation. P values indicate a one-sided t-test. **(E)** Competition assay showing that *Gm10076 KO C7* cells are strongly outcompeted by wild-type keratinocytes over 4 passages (two independent replicates). Left panels show representative images of wild-type (magenta) and *Gm10076 KO C7* (green) keratinocytes. Scale bars, 100 μm. **(F)** O-Propargyl-Puromycin (OPP) incorporation reveals a strong reduction in overall protein synthesis rates in *Gm10076 KO C7* cells compared to nontargeting control sgRNA keratinocytes. Data represent the mean of 12 OPP replicates and 3 OPP + cycloheximide (CHX) replicates, with error bars indicating the standard deviation. P values indicate a one-sided t-test. **(G)** Polysome profiles obtained through sucrose density gradient fractionation for parental Cas9::GFP, nontargeting control keratinocytes, *Gm10076 KO C7* and *Gm10076::TuGFP* overexpression keratinocytes. **(H)** P site mapping of ribosome profiling data shows strong triplet periodicity in 4 replicates of control and *Gm10076 KO C7* keratinocytes. **(I)** Ribosomal footprint length distribution in control and *Gm10076 KO C7* keratinocytes, with an expected peak at 30-31nucleotides. **(J)** *Gm10076* strongly impacts the translational landscape. Scatterplot of transcriptional and translational changes *Gm10076 KO C7* keratinocytes compared to control keratinocytes. Transcriptional changes are in blue (RNA-seq p adjusted < 0.05 and |LFC| > 0.25), homodirectional changes in orange (RNA-seq and ribosome profiling p adjusted < 0.05 and |LFC| > 0.25) and translational changes are in red (ribosome profiling p adjusted < 0.05 and |LFC| > 0.25). Histograms show the overall distribution of the transcriptional and translational changes. LFC, log_2_ fold change. **(K)** Bar graph summarizing the number of transcriptional, homodirectional and translational changes. **(L)** Gene Ontology (GO) enrichment analysis of translationally downregulated genes upon *Gm10076* perturbation. **(M)** *Rpl41* mRNA levels analyzed by RT-qPCR upon selection with either scramble or 3 different *Rpl41* shRNAs. P values indicate a one-sided t-test. **(N)** Measurement of overall protein synthesis rates via O-Propargyl-Puromycin (OPP) incorporation in wild-type and *Gm10076* KO C7 cells. Data represent the mean of 4-12 OPP replicates and 2-3 OPP + cycloheximide (CHX) replicates, with error bars indicating the standard deviation. P values indicate a one-sided t-test. **(O)** *Rpl41 shRNA; Gm10076 KO C7* cells initially exhibit stalled proliferation and require an extended adaptation period. **(P)** Representative images of *Rpl41* shRNA2; *Gm10076* KO C7 two weeks post-infection with the lentiviral *Rpl41* shRNA2 construct. Arrows mark the few surviving *Rpl41* shRNA2; *Gm10076 KO C7* keratinocytes. Scale bars, 100 μm.

To examine the function of the *Gm10076* microprotein, we established two clonal *Gm10076* knockout keratinocyte cell lines. Importantly, the sgRNAs targeting *Gm10076* do not target the canonical *Rpl41* gene, which we also confirmed by sequencing the full *Rpl41* locus (Fig. S9). Consistent with our screening data, *Gm10076* knockout cells showed reduced proliferative rates and were strongly outcompeted in competition assays with control keratinocytes (Fig. 5D-E), indicating a marked effect of *Gm10076* on proliferation.

None of the commercially available RPL41 antibodies were able to detect the protein in western blots and proteomic quantification is not feasible due to tryptic digestion producing only two amino-acid-long peptides (*47, 48*). However, our previously published Nanopore long-read sequencing data in keratinocytes and squamous cell carcinoma cells indicated that lncRNA *Gm10076* was expressed at comparable levels to *Rpl41* (Fig. S10A-B) (*49*). This raises the possibility that total RPL41 protein levels could be equally determined by both genes, *Rpl41* and *Gm10076*. The two other eukaryote-specific intersubunit bridges RPL19 and RPL24 are required for subunit joining in the context of translation initiation and elongation (*50*). We therefore hypothesized that the *Gm10076* microprotein may similarly impact intersubunit bridge formation with potential roles in initiation or elongation.

To determine the function of *Gm10076* microprotein in translation, we first tested overall protein synthesis rates in *Gm10076* knockout keratinocytes. While the *Gm10076* F7 line was identified as a heterozygous mutant with a large insertion, the C7 clonal line displayed a small homozygous 2-nucleotide deletion, implying that the otherwise wild-type lncRNA is rather unlikely to influence the phenotype (Fig. S9). Thus, our subsequent analyses focused on the *Gm10076* C7 clonal keratinocytes. Monitoring the incorporation of O-propargyl-puromycin (OP-puro) into nascent proteins, we observed a striking 40% reduction in protein synthesis rates in mutant *Gm10076* keratinocytes (Fig. 5F). Second, we analyzed polysome gradients in wild-type and *Gm10076* knockout keratinocytes by sucrose density gradient fractionation. While 40S, 60S and 80S were overall qualitatively comparable, we consistently found reduced polysomes in *Gm10076* mutant keratinocytes and higher polysome fractions upon ectopic *Gm10076* microprotein expression (Fig. 5G). In sum, these observations suggest a disruption in either efficient translation initiation or translation elongation, resulting in strongly reduced overall protein synthesis rates.

To systematically monitor how these changes in the translational machinery impact the translational landscape, we next performed ribosome profiling and matched RNA sequencing experiments in wild-type and *Gm10076* mutant keratinocytes in 4 replicates. Our ribosome profiling data set highlighted strong triplet periodicity, a feature reflecting high-quality data (Fig. 5H), as well as enrichment across the coding sequence and the expected footprint length peak at 30 nucleotides (Figs. 5I, S10C).

First, we sought to confirm our single-cell CRISPR screening data sets, which had uncovered a robust number of differentially expressed transcripts upon *Gm10076* sgRNA2 perturbation in P60 epidermal stem cells (Fig. S11A). In keeping with our single-cell CRISPR screening, we also found 2522 differentially expressed genes in *Gm10076* mutant keratinocytes by bulk RNA-sequencing (Fig. S11B, Table S4). Second, we contrasted ribosome profiling with matched RNA sequencing data, uncovering transcriptional, homodirectional and translatome-level changes in *Gm10076* knockout keratinocytes using DESeq2 (*51*). Some genes such as *Sox2, Axin2, Lox* or *Foxg1* were predominantly changed at the transcript level without concomitant alterations in translation (Fig. 5J-K). Furthermore, 1345 genes exhibited homodirectional changes with transcription and translation both significantly altered, suggesting extensive reprogramming of the gene expression landscape in *Gm10076* mutant keratinocytes (Fig. 5J-K, S12). Among the top upregulated genes in *Gm10076* mutant keratinocytes, we found an induction of *Il1a, Cxcl2, Cxcl3* and *Cxcl5*, indicating a strong inflammatory response as a consequence of cellular stress upon loss of the ribosomal microprotein RPL41 (Fig. 5J, Table S4). Finally, we also observed 456 genes that were exclusively changed at the translational level (Fig. 5J-K, S12). *Rpl41* was translationally induced, indicating a compensatory response to the loss of *Gm10076*. The *Gm10076*-dependent translationally downregulated genes were strongly enriched for the GO term translation, including various initiation factors and ribosomal genes (Fig. 5L). These observations suggest that, while *Gm10076*-deficient cells predominantly undergo secondary changes of their transcriptional programs, with subsequent homodirectional changes, the reduction of RPL41 levels selectively reduces the translational efficiency of the ribosomal machinery.

Given our findings of a second *Rpl41* in the mouse genome, we surmised that the two genes might compensate for each other, which could provide an alternative explanation for the lack of a strong phenotype upon perturbation of the conventional *Rpl41* gene. To specifically target *Rpl41* without affecting *Gm10076*, we designed shRNAs directed at the 5’UTR or 3’UTR of *Rpl41* and infected wild-type and *Gm10076* mutant keratinocytes. The 3 tested shRNAs decreased *Rpl41* mRNA levels by 45-80% in wild-type keratinocytes (Fig. 5M). Keratinocytes expressing any of the 3 *Rpl41* shRNAs were viable and showed superficially no phenotype. In line with this observation, monitoring total protein synthesis using OP-puro incorporation, we found that none of the *Rpl41* shRNAs resulted in a significant reduction of protein synthesis rates in wild-type keratinocytes (Fig. 5N). These observations suggest that the microprotein encoded on *Gm10076* can largely compensate for the knockdown of the conventional *Rpl41* gene. In contrast, although we could select the double *Rpl41* shRNA; *Gm10076* keratinocytes, the additional reduction of total RPL41 initially almost completely stalled proliferation. Proliferation remained negligible over four weeks, requiring an extended adaptation period of the *Rpl41* shRNA; *Gm10076* keratinocytes before experimental use (Fig. 5O-P, S13). Following the adaptation phase, selected double *Rpl41 shRNA; Gm10076* keratinocytes displayed only a very mild, non-significant reduction of *Rpl41* mRNA levels, in contrast to the robust knockdown efficiency observed in wild-type keratinocytes (Fig. 5M). This supports the notion that adaptation imposes selective pressure favoring cells that evade knockdown or silence the lentiviral construct, implying that *Gm10076* mutant keratinocytes cannot tolerate any further reduction in total RPL41 levels, preventing their selection. Despite this modest *Rpl41* knockdown, double *Rpl41 shRNA; Gm10076* keratinocytes showed a further significant decline in protein synthesis, with a 40% reduction upon *Rpl41 shRNA3* beyond the already strongly reduced rates observed in *Gm10076* mutants (Fig. 5N). This indicates that even the slightest additional loss of RPL41 in *Gm10076* mutants profoundly affects protein synthesis rates. Collectively, these observations strongly indicate functional redundancy between *Rpl41* and the microprotein on *Gm10076*, underscoring that the ribosome bridge protein RPL41, contrary to prior assumptions of nonessentiality, is an essential component of the ribosomal machinery.

Finally, given the functional redundancy of RPL41 and the potential adaptive control of this ribosomal bridge protein, we investigated whether RPL41 levels influence disease progression. We examined the TCGA pan-cancer atlas for ribosomal protein gene amplifications and stratified patients according to amplifications in each of the 80 ribosomal protein genes. While most ribosomal gene amplifications showed no correlation with survival, patients with *RPL41* amplification were associated with significantly reduced overall survival (Fig. S14A-C). Given the caveat of cancer type-specific confounders in this analysis, we then focused on ovarian cancer, where *RPL41* amplifications were prevalent and again found a strong significant correlation with reduced overall survival (Fig. S14D). These findings underscore the importance of maintaining balanced cellular *RPL41* levels and s uggest that excess of the ribosomal bridge protein RPL41, which could also potentially be driven by the microprotein encoded on *Gm10076*, may contribute to disease progression in cancer patients.

## Discussion

The dark proteome encompasses over 7200 microproteins that were historically neglected in genome annotations. Despite the rapidly growing number of annotated microproteins, with estimates ranging up to 50,000 (*52*), only a small fraction has been functionally characterized, raising fundamental questions about their contribution to cellular functions. Here, we established an *in vivo* single-cell CRISPR strategy to systematically investigate the tissue-wide function of microproteins at single-cell transcriptomic resolution. Our study unravels how microproteins impact cellular function in different cell types across the epidermal tissue, from stem cell populations, differentiated cells, hair follicle cells up to immune cells. We comprehensively document the global and cell-type-specific landscape of microprotein function and their impact on the cellular transcriptome, highlighting that a distinct cohort of microproteins profoundly shapes cellular gene expression programs *in vivo*.

It is noteworthy that several microproteins surfacing in our screen were officially classified as pseudogenes. These include *Gm7239* (partial alignment to *Set*) as well as the ribosomal pseudogenes *Gm10076* (full alignment to *Rpl41*), *Gm13935* (partial alignment to *Rps3a1*), *Gm5586* (partial alignment to *Rps3a1*) and *Gm8420* (pseudogene of *Rpl15*). Pseudogenes are DNA sequences with high homology to the functional gene, but have been considered dysfunctional due to the loss of regulatory sequences needed for expression. The overlap between functional microproteins in our screen and the realm of pseudogenes suggests that functional pseudogenes may be more frequent than anticipated, raising the fundamental question of how pseudogene microproteins in mammalian genomes shape the cellular proteome. The human genome contains over 2000 pseudogenes of ribosomal proteins and 22 pseudogenes for *Rpl41* (*53*). Our findings illuminate the realm of microproteins derived from pseudogenes, suggesting that they may expand our understanding of genome complexity and introduce additional layers of regulation to essential cellular machineries such as the ribosome (*54*). For instance, given that perturbation of the truncated RPS3A1 microprotein on *Gm13935* led to increased proliferation and pronounced transcriptional alterations, it is possible that this microprotein exerts an inhibitory effect by competing with RPS3A1 for ribosomal binding. As ribosomopathies, a class of human disorders driven by ribosomal protein haploinsufficiency (*55*), arise from subtle alterations in ribosomal protein levels, ribosomal pseudogenes may play a critical role in disease progression, resembling the stratification observed in cancer patients with *RPL41* amplifications (Fig. S14). Microproteins encoded from pseudogenes are therefore not necessarily nonfunctional relics but may profoundly influence cellular processes in homeostasis and disease. As such, pseudogenes may serve as an evolutionary reservoir, enabling the emergence of novel functional microproteins without necessitating *de novo* gene formation, thereby adaptively expanding the cellular proteome.

Leveraging our *in vivo* single-cell CRISPR screening strategy, we identify a second *Rpl41* gene on the lncRNA *Gm10076*. We provide multiple lines of evidence that *Gm10076* markedly contributes to total RPL41 levels, which is a microprotein located at the ribosomal intersubunit bridge. Although previously considered a nonessential ribosomal protein, our findings establish RPL41 as an essential ribosomal protein for protein synthesis and cellular proliferation and may prompt further studies on the exact role of this intersubunit bridge protein for the intricate functions of the ribosome. Our ribosome profiling data and protein synthesis analyses indicate that the *Gm10076* microprotein substantially shapes the translational landscape, likely by enabling efficient translation initiation or elongation.

Collectively, our study establishes a comprehensive framework for dissecting the functional significance of the dark proteome, uncovers a second essential *Rpl41* gene critical for ribosome function and lays the groundwork for investigating its regulation and contribution to the pathogenesis of diseases such as cancer.

## Acknowledgements

We thank Sendoel lab members for critical input on the manuscript, Catharine Aquino and the FGCZ for sequencing, the LASC Zurich for assistance with mouse work and the Cytometry Facility at UZH for assistance with sorting.

## Author contributions

F.V-F. conducted the experiments and collected the data. U.G. performed bioinformatic data processing. P.F.R. assisted with sorting and single-cell experiments. D.S. performed bioinformatic data annotation. M.O. carried out in utero lentiviral injections and animal handling. K.H. assisted with cloning. C.S. assisted with knockout cell lines and performed immunofluorescence. C.D. assisted with mouse preparations. C.D, D.T. and R.W. assisted with ribosome profiling. M.Y. assisted with OPP. M.Y. and R.W. assisted with polysome experiments. F.V-F, U.G., and A.S. performed data analysis and interpretation. A.K. and S.J.E. performed whole mount immunofluorescence staining and image analysis. H.Y. assisted with animal experiments. F.V-F. and A.S. conceived the project. A.S. supervised the project.

## Data and materials availability

The single-cell CRISPR, ribosome profiling and RNA sequencing will be available upon publication.

Valdivia-Francia et al. 2025 (preprint)

## Materials and Methods

### Data pre-processing for noncanonical ORF detection

Sequencing adapters were trimmed from raw reads using flexbar (*56*), ribosomal, mitochondrial and tRNA reads were removed by mapping against a custom genome containing only these sequences using bowtie2 (*57*). Remaining reads were mapped against the gencode GRCm38 V25M genome using the STAR aligner v2.7.7a (*58*).

### Detection of noncanonical ORFs

The Gencode GRCm38 V25M genome assembly was used for open reading frame annotation. ORF-RATER (Open Reading Frame – Regression Algorithm for Translational Evaluation of Ribosome-protected footprints) scripts were used to detect novel ORFs from published ribosome profiling data as previously described (*10, 59*) using default parameters with the following exceptions: maximum fragment length was set to 34, start codon sequence was set to “NUG” allowing for all near cognate translation initiation sites to be included, the minimum peptide length of candidates had to be 10 amino acids long and the ORF-Rater confidence score threshold was set to 0.8. Both standard CHX and harringtonine-treated samples of SCC, SOX2 and WT were used for the regression to determine the highest-confidence sites of translations.

### Candidate filtering and selection of noncanonical ORFs

Candidate files were sorted in descending order according to ORFrating (ORF-RATER score), percentQuant (perfect of nucleotides with reads used for ORF-RATER quantification), sumRibo (sum of ribosome sequencing reads across replicates) and harrReads (the sum of harringtonine reads across replicates). A total of 221 candidates were selected from the upstream (156) and new (65) categories, along with 8 known uORFs.

### Cloning of the CROPseq vector

The puromycin resistance cassette in the original CROP-seq-Guide-Puro vector (Addgene #86708) was replaced with a CRE sequence, amplified from SBE-NLS-mcherry-P2A-CreERT2 PGK-rtTA3 (Addgene #196936) and cloned in via Anza 34 Pfl23II (Invitrogen IVGN0344) and Anza 28 MluI (Invitrogen IVGN0286) restriction sites. The NLS-mcherry-P2A fragment was removed via mutagenesis Phusion DNA Polymerase reaction (Thermo Scientific, F530S).

### sgRNA design

Individual guides were designed with the Broad Institute sgRNA design tool (sgRNA Designer: CRISPRko (broadinstitute.org)) using the genomic location as input for microprotein targets. The 3 top ranking guides per target were chosen. 50 nontargeting control sgRNAs were randomly picked from the Brie library as previously described (*15*).

### Cloning of the sgRNAs into CROP-seq Cre vector

The 720 sgRNA sequences for the screen were ordered as oligo pool from IDT, cloned in batch via Gibson assembly (HiFi 1 Step Master Mix Kit, SGIDNA Synthetic Genomics GA1100-50) as previously described (*60*). For all single-cell experiments, two different library batches were cloned and individually sequenced to confirm homogeneous sgRNA representation. For the validation screen, 101 sgRNA sequences were ordered as oligo pool from IDT, cloned in batch via Gibson assembly. A single library was used for the single-cell experiment and in vitro screens and was sequenced as starting point (T0).

### Low-titer lentivirus production

Production of vesicular stomatitis virus (VSV-G) pseudotyped lentivirus was performed by calcium phosphate transfection of Lenti-X 293T cells (TaKaRa Clontech, #632180) using the construct and the helper plasmids pMD2.G and psPAX2 (Addgene plasmids #12259 and #12260), as previously described (*17*). Viral supernatant was collected 46 h after transfection and filtered through a 0.45 µm filter (Sarstedt AG, 83.1826).

### High-titer lentivirus production

High-titer vesicular stomatitis virus (VSV-G) pseudotyped lentivirus was performed by following the same procedure as for low-titer lentivirus, with an additional concentration step. The viral supernatant was concentrated ∼2,000-fold using a 100 kDa MW cut-off Millipore Centricon 70 Plus (Merck Millipore, UFC710008). The virus was further concentrated by ultracentrifugation using the SW 55 Ti rotor for the Beckman Coulter Optima L-90 Ultracentrifuge at 45,000 rpm at 4 °C for 1.5 hours. Viral particles were resuspended in viral resuspension buffer (20 mM Tris pH 8.0, 250 mM NaCl, 10 mM MgCl2, 5% sorbitol) and stored at −80 °C until used for infection.

### In vivo experiments

All animal experiments were conducted in strict accordance with the Swiss Animal Protection law and requirements of the Swiss Federal Office of Food Safety and Animal Welfare (BLV). The Animal Welfare Committee of the Canton of Zurich approved all animal protocols and experiments performed in this study (animal permits ZH074/2019, ZH196/2022).

Genetically modified mice of the strain B6J.129(B6N)-*Gt(ROSA)26Sor*^*tm1(CAG-cas9*,-EGFP)Fezh*^/J (denoted as Cas9) were purchased from the Jackson Laboratory (strain #026175). FVB.129S6-*Gt(ROSA)26Sor*^*tm1(Pik3ca*H1047R)Egan*^/J (denoted as *Pik3ca*) were originally purchased from Jackson Laboratory (strain #016977). *Cas9* were crossbred with *Pik3ca* mice to provide embryos heterozygous for the inducible Cas9 allele, suitable for lentivirus injection at embryonic development stage E9.5. Ultrasound-guided in utero injections were conducted as previously described (*61*). *Pik3ca* females at E9.5 of gestation were anesthetized with isoflurane and submitted to surgery for no longer than 30 minutes to ensure recovery. Each embryo was injected with 0.5 µl of high-titer lentivirus and up to 8 embryos were injected per litter. All mice were housed at the Laboratory Animal Services Center (LASC) of the University of Zurich in individually-ventilated cages in a humidity- and light-controlled environment (22 °C, 45–50%, 12 h light/dark cycle), and had access to food and water ad libitum. Cas9 males used for crossbreeding and pregnant Pik3ca females were housed individually. All other mice were group-housed. Successful infection of mice was controlled through excitation of the back skin with a dual fluorescent protein flashlight (NIGHTSEA, DFP-1). Positively infected mice were then euthanized by decapitation (P4) or with CO_2_ (P60), shaved and treated with hair removal cream if necessary.

P4 back skin was processed as previously described (*15*). Fat was scraped off and the back skin was washed once in cold PBS and then placed in dispase (Corning #354235) dermis-side facing down, incubated for 35 minutes at 37 °C on an orbital shaker. Next, epidermis was separated from dermis with fine forceps and placed in 4 ml 0.25% Trypsin-EDTA (1×, Gibco; 25200056) for 15 minutes at 37 °C with orbital shaking. Epidermis was torn into smaller pieces and washed with cold PBS, pipetted vigorously up and down to achieve a single-cell suspension. Suspension was filtered through a 70-µm and subsequently a 40-µm strainer (Corning; #431750, #431751). Suspension was then centrifuged for 10 minutes at 400 *g* and resuspended in FACS buffer (PBS + 2% chelexed FBS (FBS(-)) + DAPI).

P60 back skin was processed as previously described (*15*). Back skin was harvested with surgical scissors and placed dermis-facing up on a styrofoam rack with pins. Fat was scraped off with a scalpel. Fat-free skin was then washed, dermis-side facing down in 1× PBS. Skin was then placed in the same orientation in 0.5% Trypsin-EDTA (10×, Gibco; 15400054) and incubated at 37 °C on an orbital shaker for 25 minutes (females) or 50 minutes (males). Using a glass microscopy slide, skin was secured to the bottom of the dish and scraped again with a scalpel until it started to diverge. Excess cold PBS with 2% FBS(-) was added to neutralize trypsin. Cell suspension was strained on ice first through a 70-µm strainer, washed with an additional 15 ml of cold 1× PBS with 2% FBS (-) and then filtered through a 40-µm strainer. Cell suspension was spun down at 400 *g* for 10 minutes and resuspended in FACS buffer.

### Single-cell RNA sequencing

EGFP-positive cells were sorted on a BD FACSAria III using a 70-µm nozzle and processed for single-cell capture with a BD Rhapsody single-cell analysis system using BD Rhapsody cartridge kit (#633733). For P4, 7 separate cartridges with an average of 76,613 cells were loaded. For P60, 8 cartridges with an average of 68,171 cells were loaded. Reverse transcription from beads and sequencing library production was carried out according to the manufacturer’s instructions (BD Biosciences, doc ID: 210967 rev. 1.0).

P4: a total of 18 injected P4 mice were collected and assessed in 7 independent single-cell RNA sequencing runs.

P60: a total of 7 injected mice were collected and assessed in 8 independent single-cell RNA sequencing runs.

### Dial-out nested PCRs for sgRNA amplification

As previously described (*15*), the sgRNA-containing region was amplified from the BD Rhapsody beads using a nested KAPA Polymerase reaction. In the first polymerase reaction (PCR1), the Rhapsody beads from each separate cartridges were resuspended and distributed into 4 separate 50 µL reactions (100 µl KAPA HiFi HotStart ReadyMix (Roche 07958935001), 6 µl forward primer (5′-ACACGACGCTCTTCCGATCT-3′, 10 µM), 6 µl reverse primer (5′-TCTTGTGGAAAGGACGA-3′, 10 µM), 12 µl Bead RT/PCR Enhancer reagent from BD Biosciences and 72 µl nuclease-free water for a total volume of 200 µl). PCR1 conditions were as follows: initial denaturation at 95 °C for 5 min, followed by 25 cycles of denaturation 95 °C for 30 s, annealing 53 °C for 30 s, extension 72 °C for 20 s and a final extension of 10 min at 72 °C.

The 4 × 50 µl PCR1 product was pooled, and beads were removed using a magnet. Amplicons were cleaned up using Agencourt AMPure XP beads (Beckman Coulter, A63881) following manufacturer’s instructions. 3 µl of cleaned up PCR1 was used as template for PCR 2 (25 µL KAPA HiFi HotStart Ready Mix, 2 µl forward primer (5′-ACACGACGCTCTTCCGATCT-3′, 10 µM), 2 µl reverse primer (CAGACGTGTGCTCTTCCGATCTCTTGTGGAAAGGACGAAACA*C*C* G-3′, 10 µM), 18 µl nuclease-free water) for a total volume of 50 µl. PCR2 conditions were as follows: initial denaturation at 95 °C for 3 min, followed by 10 cycles of denaturation 95 °C for 30 s, annealing 60 °C for 3 min, extension 72 °C for 60 s, and a final extension of 5 min at 72 °C. Agencourt AMPure XP beads were used to clean up PCR2 product according to the manufacturer’s instructions before the indexing PCR. Indexing PCR was performed according to BD Biosciences ‘mRNA Targeted Library Preparation’ protocol (doc ID: 210968 rev. 3.0).

### Sequencing of libraries

All single-cell or amplicon sequencing for the P4 and P60 timepoints and the initial T0 libraries were prepared as ready-made libraries and sequenced either on Illumina NovaSeq or Nextseq 500 instruments at the Functional Genomics Center Zurich. Prior to sequencing, libraries were checked for peak size and concentration on a Bioanalyzer 2100 using DNA High Sensitivity Chip (Agilent, 5067-46). Concentrations were additionally measured using a Qubit 1X dsDNA High Sensitivity Assay (Invitrogen, Q33230) and libraries were prepared with equal representation of each whole-transcriptome amplification (WTA) and dial-out amplicon. WTA amplification analyses from Rhapsody Cartridges were sequenced in full SP flow cells at 1.8 nM concentration with 20% PhiX and the following read configuration: Read 1 60, index 1 8 cycles, read 2 62 cycles.

### Primary keratinocyte isolation and culture

Primary mouse epidermal keratinocytes were isolated from 4-day old C57BL/6 wild-type mice, *Cas9* mice and *Pik3ca* mice as previously described (*62*). Briefly, isolated epidermal keratinocytes were cultured on 3T3-S2 feeder layer pre-treated with Mitomycin-C, using 0.05 mM Ca^2+^ E-media supplemented with 15% serum. After 3 passages on 3T3-S2 feeder layer, cells were transferred to 0.05 mM Ca^2+^ E-media, prepared in-house, as previously described (*63*). The C57BL/6 wild-type keratinocytes were used for lentiviral infection and protein sample collection and immunofluorescence. All cell cultures were maintained under standard conditions at 37°C and 5% CO_2_.

### *In vitro* lentivirus infections

Lentiviral infections for the generation of knockout cell lines were performed using *Cas9* keratinocytes. Cells were seeded at a density of 1.5-2.5 × 10^5^ cells per well in a 6-well plate (Thermo Scientific Nunclon TM Delta Surface; #140675). Cells were infected with 100-300 µl of low-titer virus in the presence of the infection mix (1/10 dilution of polybrene [10 mg/ml Sigma; 107689-100MG in PBS] in FBS[-]), followed by centrifugation of the plates at 1100 *g* for 30 minutes at 37 °C in a Thermo Heraeus Megafuge 40R centrifuge. Infected cells were FACS-sorted with BD FACSAria III using a 70 µm nozzle. For the generation of overexpression cell lines, a similar protocol was followed using C57BL/6 wild-type keratinocytes.

### Cloning individual sgRNAs for guide efficiency

45 sgRNAs representing top, middle and bottom enrichment cohorts were selected and sgRNA efficiency for was assessed by infecting *Cas9* keratinocytes. Low-titer lentiviral preparations of the CROP-Seq-Puro vector with individual sgRNAs was produced, cells were infected and selected with puromycin (1 µg/mL, Gibco; A11138-03). Cells were harvested after 10 days of selection. Genomic DNA was extracted using a DNeasy Blood and Tissue Kit (QIAGEN; 69504), followed by Phusion DNA Polymerase reaction (Thermo Scientific, F530S) according to manufacturer’s instructions, using GC-rich 5X buffer. Amplicons were 600-1000 bp long and amplified by primers flanking the sgRNA. Primers were designed with primer 3. TIDE analysis was performed to evaluate the edits.

### Cloning shRNAs for *Rpl41* knockdown

To knock down *Rpl41* in nontargeting and *Gm10076* C7 mutant keratinocytes, 3 different shRNAs were cloned into a modified LV RFP plasmid (Addgene #26001), where H2B-RFP was replaced with a blasticidin selection marker. First, the Pfl23II restriction site was inserted before H2B RFP via a Phusion mutagenesis Polymerase reaction (Thermo Scientific, F530S). The blasticidin selection cassette was amplified from C1(1-29)-TurboID-V5_pLX304 plasmid (Addgene # 107175) with primers containing Pfl23II restriction site overhang in the forward primer and KpnI restriction site overhang in the reverse primer. shRNAs were inseted via Phusion DNA Polymerase reaction (Thermo Scientific, F530S) using the following sequences: scramble shRNA: CAACAAGATGAAGAGCACCAACTCGAGTT-GGTGCTCTTCATCTTGTTGTTTTT

*Rpl41* shRNA1: CATCGGTAATGAGTCTCAGTATTCAAGAGATACTGAGACTCATTACCGATGTTTTT

*Rpl41* shRNA2: ACTGTGTGCTGCCATCGGTAATTCAAGAGATTAC-CGATGGCAGCACACAGTTTTTT

Rpl41 shRNA3: GCACGCCATTAAATAGCAGTATTCAAGAGATACTGCTATTTAATGGCGTGCTTTTT

### Overexpression constructs

For overexpression constructs with microproteins *Gm13935, Gm10076, Gm5586, Gm8420* and *Gm7239*, gBlocks (IDT) were ordered with the nucleotide sequence of the microproteins. At the 5’end of the microprotein sequence (Table S5), a XbaI restriction site was added. A linker sequence GSPGS followed by the Pfl23II restriction site was added at the 3’ end of the microprotein. 100 ng of microprotein gBlocks were digested and ligated into a pLKO TuGFP plasmid using the Anza 12 XbaI (Thermo scientific, IVGN0126) and Anza 34 Pfl23II (Thermo scientific, IVGN0344) restriction enzymes.

### Western blot

Keratinocytes and Hek293T cells were collected from a 10 cm culture plate when they reached 80-90% confluency. Cells were washed with cold 1x PBS (Gibco 10010-015), scraped off the plate and lyzed for 10 minutes on ice with RIPA buffer (Sigma R0278) with 1X Protein Cocktail inhibitor (50X Promega, G6521). Protein lysates were quantified using Pierce BCA Protein Assay Kit (Thermo Scientific #23225) according to manufacturer guidelines. 20 µg of protein lysate was mixed with NuPAGE LDS Sample buffer (4X, Invitrogen NP0007) and NuPAGE Sample Reducing Agent (10X, Invitrogen NP0009). Samples were heated to 95°C for 5 minutes and loaded on a 4-12% NuPage 12-well Bis-Tris Gel (Invitrogen, #7001691) using 1X NuPAGE MOPS SDS Running Buffer (20X, Invitrogen, NP0001). Subsequently, proteins were transferred to a nitrocellulose membrane (Cytiva Amersham Protran 0.45 µm NC, 10600002) by tank transfer at 4°C, 30V for 90 minutes. Membranes were blocked in 5% skim milk for 2 hours at room temperature. Primary antibodies were incubated overnight at 4°C, followed by 3 washes with TBS-Tween 0.1%. Secondary antibodies were incubated for 2 hours at 4 °C followed by 3 washes with TBS-Tween 0.1% before developing with freshly mixed ECL solutions (Amersham Cytiva. RPN2209). The following antibodies were used: Anti-turboGFP Rabbit polyclonal antibody (ORIGENE TA150071), PTEN (138G6) rabbit mAB (Cell signaling #9559), Actin (8H10D10) mAB (Thermo Scientific MA5-15452), Cas9 (7A9-3A3) mouse mAb (Cell signaling #1497), Anti-rabbit IgG HRP-linked antibody (Cell signaling #7074) and Anti-mouse IgG HRP-linked antibody (Cell signaling #7076). Western blot bands were quantified using ImageJ.

### Immunofluorescence

Fixed back skin pieces cut as strips along the AP axis were embedded in OCT (Tissue-Tek; #4583) and frozen over dry ice. 10 µm thick sagittal sections were obtained using a Micron HM550 OM cryostat (Thermo Scientific) and immobilized on Superfrost glass slides. Slides were washed twice in 1× PBS before blocking in blocking solution (1% BSA, 1% gelatin, 5% normal donkey serum, 0.30% Triton X-100, 1× PBS). Primary antibodies CD324 (E-Cadherin) anti-rat antibody (Invitrogen, 14-3249-82) and 1:200, GFP anti-chicken antibody (Abcam, ab13970), 1:200 were incubated overnight at 4 °C. After washing twice with 1× PBS, slides were incubated with secondary antibodies (Alexafluor488 and Cy3, Jackson ImmunoResearch Laboratory; 1:500) and 1 µg/ml 4′,6-diamidino-2-phenylindole (DAPI) at room temperature for 1 h. Sections were then washed again with 1× PBS, dried, covered with ProLong™ Diamond Antifade Mountant (invitrogen, P36965) and sealed with a coverslip and allowed to cure for 24 hours before imaging.

### Wholemount immunofluorescence and antibodies

Dissected back skin, injected with the library, was fixed in 4% PFA for 1 h at room temperature. Following fixation, samples were permeabilized in 1X PBS containing 1% TritonX-100 overnight. All steps during staining were carried out at room temperature. Primary antibodies were diluted into blocking buffer (5% donkey serum, 2.5% fish gelatin, 1% BSA, 1% Triton in PBS, all antibodies diluted 1:200) and were incubated for at least 16–20 h at room temperature. Samples were then washed for 3–4 h in 1% PBS-Triton, and incubated with secondary antibodies (in blocking buffer, 1:200) together with DAPI (1 µg/ml) for at least 16–20 hours. After staining, samples were extensively washed with 1% PBS-Triton every hour for 4–6 h. Samples were then dehydrated in increasing concentrations of Ethanol: 30%, 50% and 70% in doubled-distilled water (with the pH adjusted to 9.0 with NaOH/HCL) for 1 h each, and finally in 100% ethanol for 1 h. For tissue clearing, samples were transferred to Eppendorf tubes with 500 µl ethyl cinnamate (ECi, Sigma 112372) and shaken overnight at room temperature under dark conditions. Fresh ECi was used for mounting for imaging.

### Imaging and image processing

Tiled images of sectioned tissue were acquired on a Zeiss LSM900 microscope using an LD LCI Plan-Apochromat 40x/1.2 Multi-Immersion, WD 0.41 mm objective with glycerine as the immersion medium. Images were stitched using Zen Blue (Zeiss).

Tiled images of wholemount samples were acquired using a Visitron Spinning Disk with a Yokogawa CSU-W1-T2 spinning disk (50 µm pinholes, spacing 500 µm, 4000 rpm), a CFI Plan Apo Lambda 20x/0.75, working distance (WD) 1.00 mm objective and an EM-CCD camera. Images were stitched using Visiview (Visitron Systems). Further processing for all images was done in Fiji (ImageJ). All images depicted are maximum intensity projections.

### MG132 treatment

Keratinocytes expressing the constructs *Gm13935::TuGFP, Gm13935::V5P2A::TuGFP, Gm13935::P2A::TuGFP* and *TuGFP::V5::Gm13935*, were seeded at a density of 120,000 cells in a 6-well plate. 24 hours later, cells were treated with MG132 (Merck, M7449-200UL) at a final concentration of 2 µM or 10 µM for 2 hours. As control, cells were treated with DMSO. After 2-hour treatment, cells were lyzed with RIPA buffer and collected for western blot analysis as previously described in the Western blot section. Similarly, cells were seeded in glass coverslips and processed for immunofluorescence to check the expression of turbo GFP. For a time course experiment, *Gm13935::TuGFP, TuGFP::V5::Gm13935* expressing keratinocytes and wild-type keratinocytes were seeded at a density of 120,000 cells in a 6-well plate. 24 hours later, they were treated with 2 µM of MG132 for 2 hours, 4 hours and 8 hours. Treatment was started sequentially, and all samples were collected at the end of the 8-hour time point for western blot analysis as previously described.

### Overexpression constructs

For overexpression constructs with microproteins *Gm13935, Gm10076, Gm5586, Gm8420* and *Gm7239*, gBlocks (IDT) were ordered with the nucleotide sequence of the microproteins. At the 5’end of the microprotein sequence (Table S5), a XbaI restriction site was added. A linker sequence GSPGS followed by the Pfl23II restriction site was added at the 3’ end of the microprotein. 100 ng of microprotein gBlocks were digested and ligated into a pLKO TuGFP plasmid using the Anza 12 XbaI (Thermo scientific, IVGN0126) and Anza 34 Pfl23II (Thermo scientific, IVGN0344) restriction enzymes.

### Immunofluorescence of overexpression constructs and MG132 treated cells

Previously infected keratinocytes with microprotein constructs were seeded at a density of 5×104 cells on glass coverslips in a 24-well plate to reach a 70% confluency. Cells were washed and processed for immunofluorescence 24 hours post-seeding. Cells were fixed with 4% Paraformaldehyde (16% Formaldehyde solution (w/v), Methanol-Free, Thermo Scientific 28908) in PBS for 15 minutes, permeabilized with 0.3% Triton X-100 in PBS for 15 minutes at room temperature, and blocked with blocking buffer (1% BSA, 1% Gelatin fish skin, 5% Serum, 0.3% TritonX-100 in PBS) for 1 h at room temperature. Cells were stained with primary antibody overnight at 4 °C, followed by 3 washes with PBST (0.1% TritonX-100 in PBS). Secondary antibody was subsequently applied with DAPI in blocking buffer for 45 minutes at room temperature in the dark, followed by 3 washes with PBST. Cells were mounted with homemade mounting media and imaging was performed on Zeiss Axio Observer 7. The following antibodies were used: Anti-turboGFP Rabbit polyclonal antibody (ORIGENE TA150071), AF488 (Jackson Immuno Research 712-545-153).

### MTT assay

Proliferation rates for *Gm10076* knockout cell lines and control cell lines were tested using the MTT Cell Growth Assay Kit (Merck CT01). Keratinocytes were seeded at a density of 15,000 cells per well in a flat bottom 96 well plate, 24 hours prior to the assay. 10 µL of freshly made MTT was added to each well and incubated for 4h at 37°C, 5% CO2 to allow for the production MTT formazan. Crystals were dissolved by adding 100 µL of isopropanol with 0.4N HCl to each well with vigorous pipetting. Within 1 hour, the absorbance was measured with a test wavelength of 570 nm and a reference wavelength of 630 nm using a Tecan Infinite M1000 Pro Plate reader.

### Competition assay

To determine the rate of proliferation, nontargeting control keratinocytes and *Gm10076* C7 knockout cells were seeded at an equal density (100,000 cells each) in a 6-well plate. Percentage of mCherry-positive cells (nontargeting control) and GFP-positive (*Gm10076 C7*) were measured every passage for a total of 4 passages using Flow Cytometer Analyser BD Fortessa. In addition, wells were imaged under the Zeiss Axio Observer 7 microscope after every passage.

### Sucrose gradient density fractionation

Keratinocytes were seeded into 15 cm plates 24 hours prior to collection. The following day, 90% of the media was removed and fresh media was added 3 hours prior to harvest. Cells were allowed to reach 85-90% confluency. To separate polysome fractions in wild-type and Gm10076 C7 clonal keratinocytes, lysates were prepared using a lysis buffer (20 mM Tris-HCl (pH 7.4), 150 mM NaCl, 5 mM MgCl_2_, 1% Triton X-100, 0.5% NP-40, 1 Mm DTT, and 100 µg/ml cycloheximide). Polysomes were then fractionated on a 10–50% sucrose gradient prepared in gradient buffer (20 mM Tris-HCl pH 7.4, 150 mM NaCl, 5 mM MgCl_2_, 100 µg/ml cycloheximide) by ultracentrifugation at 41,000 rpm for 1 hour 50 minutes. The Biocomp Density Gradient Fractionation System was used to collect polysome fractions.

### Processing of ribosome profiling samples

Nontargeting control and *Gm10076 C7* keratinocytes were seeded into a 15 cm plate 24 hours prior to collection. At 80% confluency the following day, 90% of the media was removed and fresh media was added 3 hours prior harvest. Cells were washed once with 10 mL PBS CHX (20 µg/mL). PBS CHX was removed, and plates were flash-frozen by letting them float in liquid nitrogen. 400 µL of lysis buffer (1x mammalian polysome buffer, 1mM DTT, 1% TritonX-100, 0.5%NP40, 25U/µL Turbo DNase, 100µg/mL CHX) was added to the plate. When plates were partially thawed, cells were scraped to collect them into a RNase-free tube. Samples were incubated for 10 minutes on ice and centrifuged at 20000 *g* for 10 minutes at 4°C. Supernatant was transferred to a new tube. RNA concentration was measured using a Qubit RNA broad range kit. Samples were snap-frozen in liquid nitrogen and stored at −80°C until further use. Four individual replicates were collected for each condition.

### Ribosome profiling

Ribosome profiling was conducted with some adjustments of the original method (*64–66*). For RNA digestion, cell lysates were incubated with 1 U/µg of RNase 1 (Epicentre/Lucigen, 10U/µL) for 45 minutes at room temperature. Reaction was stopped by adding 4 µL of SUPERase Inhibitor (20 U/µL, SUPERase-In RNase Inhibitor Thermo Scientific AM2694). To eliminate free nucleotides and isolate ribosome-protected fragments, the RNase I-treated samples were processed using Amersham MicroSpin S-400 HR columns (Cytiva, 27514001). RNA extraction was performed utilizing TRIzol LS Reagent (Invitrogen 10296028) in combination with the Directzol RNA MiniPrep kit (Zymo Research, R2052). Total RNA concentration was measured using Qubit RNA broad range kit (Invitrogen, Q10210). Ribosome footprints were excised and extracted from a 15% TBE-UREA gel (Invitrogen, EC68852BOX) using the 20nt, 22nt, 27nt, 30nt and 60nt oligonucleotides mix as markers (1 µL of 10 µM dilution in 19µL RNase-free H2O). Extraction was performed overnight with RNA extraction buffer (300 mM NaOAc pH=5.5, 1mM EDTA, 0.25% v/v SDS). T4 PNK (NEB, M0201S) end-healing reaction was performed on the ribosome footprints for 1 hour at 37°C. Linker ligation reaction followed using T4 RnI truncated K227Q (NEB, M0351L) in combination with 20 µM of preadenylated linker for 3 hours at 22°C (50% w/v PEG-8000, 10X T4 RNA ligase, preadenylated linker, T4 RnI truncated K227Q). Ribosomal RNA was depleted using biotinylated rRNA depletion oligos (2 µM) by mixing 10 µL of sample with 10 µL of oligo mix and incubated for 2 minutes at 80°C, followed by a gradual decrease in temperature to 25°C. MyOne Streptavidin dynabeads (Life Technologies, 65001) were used to recover the samples. Reverse transcription was performed on the Ribosomal RNA-free samples using SuperScript IV Reverse Transcriptase (Invitrogen, 18090050). Footprints were excised and extracted from a 15% TBE-UREA gel. Extraction was performed overnight with DNA extraction buffer (10mM Tris-HCl pH=8, 300mM NaCl, 1mM EDTA). cDNA was circularized using CircLigase II (100 U/µL, Biosearch, CL4115K) for 1 hour at 60°C. cDNA was amplified using Phusion High-Fidelity DNA polymerase (Thermo Scientific, F530S) using an individual index per sample. 6-12 cycle amplicons were visualized using an 8% TBE non-denaturing polyacrylamide gel to determine optimal cycle number. PCR was carried out with 7-9 cycles and analyzed on an 8% TBE non-denaturing polyacrylamide gel (Invitrogen, EC61252BOX). DNA was excised from gel and extracted using DNA extraction buffer by incubating overnight with gentle shaking. DNA was recovered and precipitated in 10 mM Tris-HCl pH=8. DNA was quantified using Qubit 1X dsDNA High Sensitivity Assay (Invitrogen, Q33230) and quality was verified using the Bioanalyzer High Sensitivity DNA kit (Agilent, 5067-4626). Samples were pooled and prepared for sequencing.

### Bulk RNA sequencing

RNA was isolated from lysates from nontargeting control and *Gm10076 C7* keratinocytes as described in the Ribosome profiling section using TRIzol LS Reagent (Invitrogen, 10296028) and column purified with a Direct-zol RNA MiniPrep kit (Zymo Research, R2052). Libraries were prepared and sequenced at Novogene.

### RT-qPCR

*In vitro* Gm10076 C7 KO and nontargeting control mCherry keratinocytes, infected with pLKO Blasticidin scramble shRNA, *Rpl41* shRNA1, *Rpl41* shRNA2 and *Rpl41* shRNA3 were lyzed in 6-wells with 960µL TRIzol Reagent and the RNA was purified using the Direct-zo RNA MiniPrep kit (Zymo Research). The procedure was performed according to the manufacturer protocol except for an additional 1 minute centrifugation after the last washing step to completely remove potential ethanol leftover. RNA was eluted in 12 RNA/DNA free water and 1 µL was used to measure the concentration using the Qubit RNA BR Assay Kit (Invitrogen Q10210). cDNA was synthesized using the GoScript Reverse Transcription Mix, Oligo (dT) (Promega A2791). Following manufacturer protocol, 500 ng of RNA were converted into oligo (dT)-primed first-strand cDNA. iTaq Universal SYBR Green Supermix was used according to the manufacturer’s protocol for RT-qPCR reaction in a QuantStudio 7 Flex (Applied Biosystem by Life Technologies). qRT-PCR primers for *Rpl41* f: CACCGAGCACGCCATTAAATAG r: CTTGGACCTCTGCCTCATCTTT and for Gapdh f: AGGTCGGTGTGAACGGATTTG and r: TGTAGACCATGTAGTTGAGGTCA. The delta-ct method in Quant Studio Real-Time PCR software (v1.3) was used for analysis and to calculate fold changes based on ct values.

### Protein synthesis analysis

For overall protein synthesis analysis, O-Propargyl-puromycin (OPP, MedChem, MCE-HY-15680) was administered to the different cell lines 1 hour before fixation, followed by staining with Alexa Fluor 647 azide ClickiT reaction (VF 647 Click-iT EdU Universal Cell Proliferation Detection Kit, MedChem, HY-K1084). As a control, translational elongation inhibitor cycloheximide was administered 30 minutes prior to OPP administration at a final concentration of 100 µg/mL. In short, 1.2-1.5 x10^5^ keratinocytes were seeded on 12-well plates (Thermo Scientific Nunclon TM Delta Surface; 140675) 24 hours prior to the experiment. Media was changed 3 hours before the cycloheximide treatment and OPP treatment. After OPP treatment, cells were washed once with PBS and 300 µL of 0.25%Trypsin-EDTA was used to collect the cells. Cells were incubated with 200 µL of 4% PFA for 30 minutes at room temperature. Cells were incubated 5 minutes with glycine (2 mg/mL) and washed with PBS 3% BSA for 5 minutes. Cells were then permeabilized using PBS 1% TitronX-100 for 10 minutes, followed by a wash with PBS 3% BSA for 5 minutes. Cells were then incubated with Click reaction using Alexa Fluor 647 azide for 30 minutes in the dark. Cells were then resuspended in PBS 1% BSA and analyzed via Flow Cytometer on a BD Fortessa.

### Bioinformatic analysis of single-cell RNA sequencing data

Raw sequencing data, consisting of BCL files, were demultiplexed using Illumina bcl2fastq v2.20 with default settings. One mismatch in the sample barcode sequences was allowed. The resulting fastq files were processed using the BD Rhapsody Pipeline (version 1.9.1) hosted at the Seven Bridges’ cloud platform (https://www.sevenbridges.com/). A custom mouse reference genome, based on Gencode vM25 (GRCm38.p6), was generated and combined with the 720 sgRNAs sequences. Based on STAR aligner, a custom STAR reference genome was created and the Refined Putative Cell Calling option was disabled during the pipeline execution. For downstream analysis, unique molecular identifier (UMI) count data was corrected using BD Genomics Recursive Substitution Error Correction (RSEC).

### Processing of sgRNA dial-out data

PCR dial-out data was processed using an in-house custom Python script as previously described (*15*). Three 9-nucleotide cell barcodes were extracted from the 97 possible barcodes from Read 1. One mismatch was allowed per barcode (Hamming distance of 1). The script also retrieved the 8-nucleotide UMI sequences from Read 1, as specified in the BD manual. For valid barcodes, sgRNAs were identified in Read 2 by performing an exact match search for the 20-nucleotide sgRNA sequence. UMI counts per cell were deduplicated and sgRNA UMI counts per cell were computed. If multiple sgRNAs were detected for one cell, the cell was assigned to a specific sgRNA only if its UMI count exceeded the 99^th^ percentile.

### Processing of single-cell RNA sequencing data

The Scrublet Python package was used to remove doublets from each dataset separately. Genes expressed in a minimum of 5 cells were retained and an expected double rate of 20% (as estimated by BD Rhapsody) was applied. After filtering, seven P4 single-cell RNA-seq datasets and eight P60 datasets were merged using Seurat(*67*). Only cells containing detected sgRNAs and classified as singlets were selected for further analysis. Cells were filtered based on the following criteria: UMI counts > 500, UMI counts lower than the 0.99 quantile (to remove outliers) and mitochondrial gene content < 20%. Filtered datasets were then processed using Seurat’s standard pipeline. Gene expression was normalized using the NormalizeData function, which scales counts to 10,000 per cell followed by a natural log transformation (log1p). The FindVariableGenes function identified 2,000 highly variable genes, and data scaling was performed using ScaleData. Principal Component Analysis (PCA) was conducted with RunPCA and the ElbowPlot function was used to determine the top principal components that contributed to explaining the variance of the data for each dataset at P4 and P60.

The FindNeighbors function was applied to identify nearest neighbors based in the selected principal components. Clustering was performed using FindClusters with Louvain modularity optimization. To visualize the data in two dimensions, UMAP (Uniform Manifold Approximation and Projection) was used. No batch effects were observed in the P4 and P60 datasets, so batch effect correction was not performed.

To annotate cell clusters, FindAllMarkers was used to identify marker genes, and clusters were assigned to cell types based on the expression of known marker genes. Differential expression analysis was conducted using the MAST package and the Wilcoxon rank-sum test, which produced comparable results. Additionally, kernel density estimates for gene expression were inferred using the Nebulosa package.

### sgRNA enrichment and depletion calculation

The number of sgRNAs in the amplicon sequencing data were counted by the count command of the MAGeCK package (*68*). To calculate enrichment and depletion of sgRNAs, we used amplicon counts detected at P4 and P60 compared to the sequenced pre-injection libraries (A and B) originating from the different library batch preparations. The test command of the MAGeCK package was used for comparing P60 vs T0, P4 vs T0 and P60 vs T0 with control sgRNAs used for normalization and for generating the null distribution of RRA.

### Differential expression analysis

For single-cell differential expression analysis, the glmGamPoi R package (*69*) was used, which employs a Gamma-Poisson generalized linear model to the data. Genes that were expressed in at least 10% of the cells were selected, similar to the Seurat differential expression approach. The Gamma-Poisson generalized model was applied to our data using the standard settings of the glm_gp function. A quasi-likelihood ratio test for a Gamma-Poisson fit was performed with the test_de function in order to obtain the P-values for DE. The P-alues were corrected for false discovery rate using the Benjamini-Hochsberg procedure.

RNA-seq and Ribo-seq DE and TE analysis were performed by DESeq2. The normalization factors for each sample were calculated by the estimateSizeFactors function using the median of ratios method. Then the estimation of dispersion was conducted by the estimateDispersions function.

Next, a negative binomial generalized linear model was fitted for each gene to identify differentially expressed (DE) genes. The p values were obtained by the Wald test and were corrected for false discovery rate using the Benjamini-Hochsberg procedure.

### Bulk RNA sequencing data analysis

The Nextflow RNA-seq pipeline (https://github.com/nf-core/rnaseq) version 3.17.0 was used to process the paired-end RNA-seq data. Briefly, the quality of the fastq files was checked by the FastQC program followed by quality and adapter trimming by TrimGalore. The reads were aligned to the Gencode vM25 (GRCm38.p6) reference by STAR. Transcriptome quantification was performed by Salmon. MultiQC was used for the quality control of analysis pipelines.

### Ribosome profiling data analysis

A custom “regular expression pattern” together with UMI-tools was used to extract ribosome footprints from the fastq files. Footprints with minimal length of 20 nt were extracted, along with a valid sample barcode of 5 nt length. Adapter sequences AGATCGGAAG were moved to the 7 nt UMI sequence of the fastq header for further processing. Next, the reads aligning to rRNAs and tRNAs were removed from the sequencing data using bowtie2 v2.2.5. The remaining reads were aligned to the Gencode vM25 (GRCm38.p6) reference genome by STAR v2.7.11a allowing one mismatch (--outFilterMismatchNmax 1) and max 20 multimappers (--outFilterMultimapNmax 20). The UMI deduplication of the alignment bam files was performed by the UMI-tools dedup command. The reads mapping to the CDS regions of the genome in forward strand were counted by featureCounts program.

Ribowaltz R package was used for Ribo-seq quality control and periodicity analysis (*70*). The deduplicated transcriptome bam files were loaded into Ribowaltz by bamtolist command using M25 gencode annotations for protein coding transcripts. The psite offsets were computed by the psite command in auto mode with flanking 6. Next, the periodicity and other QC plots were produced from different Ribowaltz functions.

### Pathway enrichment

Pathway enrichment analysis was performed by EnrichR (https://maayanlab.cloud/Enrichr), which uses a combination of statistical methods including Fisher’s exact test, Z-score and combined score to assess, MSigDB Hallmark 2020, GO Biological Process and other gene sets. A cut-off of FDR < 0.05 was determined to select differentially expressed genes. We also conducted pathway enrichment analysis using the GSEA preranked method using GSEApy (*71*) for the analysis of up- and downregulated genes, improving the sensitivity of the gene set enrichment analysis.

### Perturbation-perturbation matrix

To characterize the transcriptional phenotype of each perturbation, we calculated the average gene expression for each perturbation in epidermal stem cells at P60. The top 2000 most variable genes across perturbations were selected and a correlation matrix was generated. Heatmaps were constructed using the pheatmap package and clustering was performed using the ward D2 method. We used the preserve neighbors from the PyMDE package to generate a minimum-distortion embedding of the data in two dimensions.

### General data analysis

Analysis was carried out using in-house Python 3.9 and R 4.1 scripts. Data wrangling was performed with the pandas library in Python and tidyverse library in R. Heatmaps were generated using the ComplexHeatmap circlize and pheatmap packages. Other plots were made using the ggplot2 library in R and seaborn library in Python.

**Supplementary Fig. 1:**
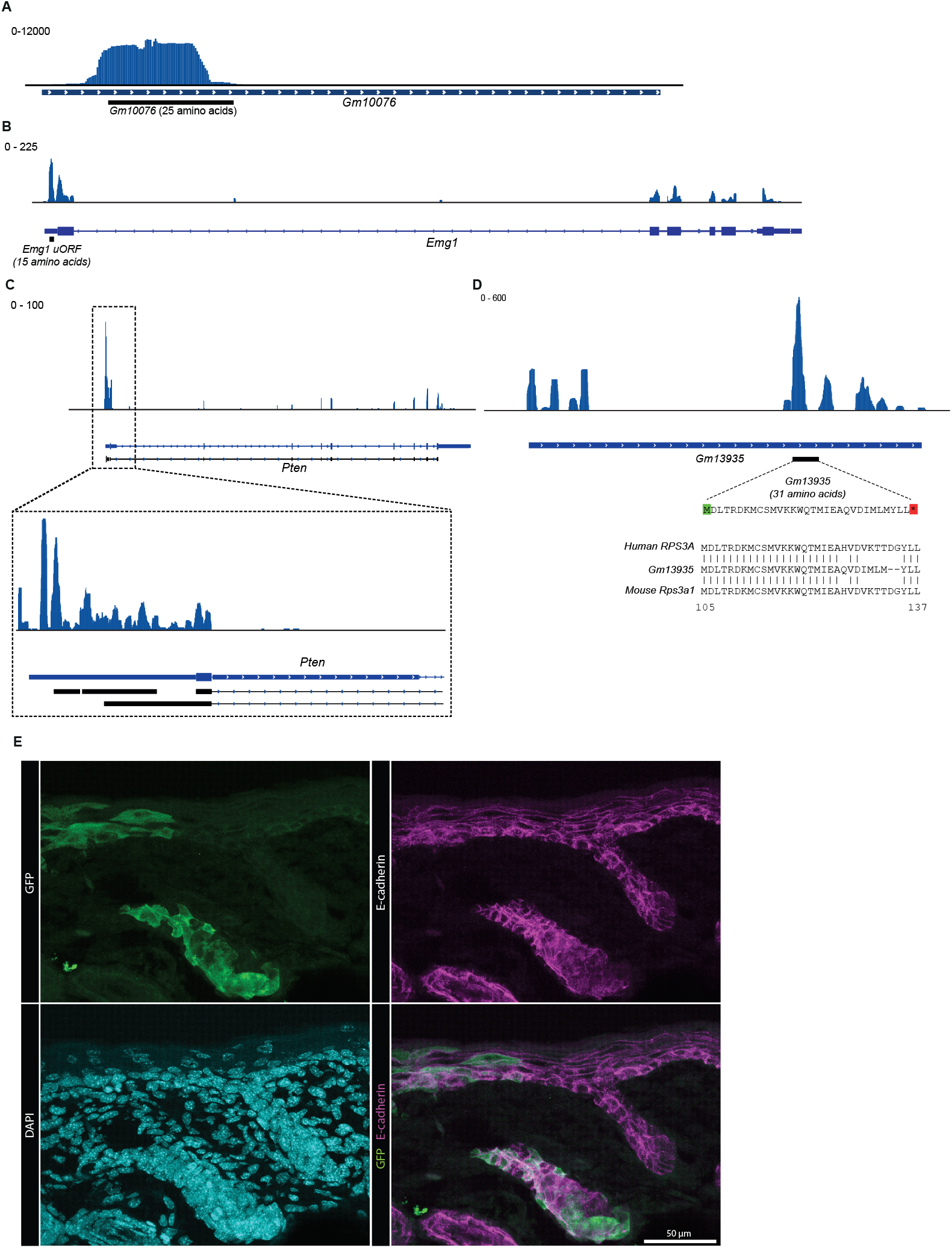
Ribosome profiling identifies microproteins translated from lncRNAs and 5’UTRs. **(A)** Ribosome footprints identify translation of a 25 amino acid microprotein on the lncRNA of *Gm10076* in Chromosome 14. **(B)** Ribosome footprints identify a 15 amino acid microprotein on the 5’UTR of the *Emg1* gene on Chromosome 6. **(C)** Ribosome footprints identify two uORFs in the 5’UTR of *Pten*. The first uORF is a 45 amino acid microprotein in frame with *Pten* CDS. The second uORF, located in closer proximity to the *Pten* CDS, is an out of frame 128 amino acid microprotein. **(D)** Ribosome profiling identifies a 31 amino acid microprotein on the lncRNA of *Gm13935*. This 31 amino acid microprotein aligns to *Rps3a1*, spanning amino acids 105 to 137. **(E)** Higher magnification of a GFP-positive clone in a section of the P4 epidermis. E-cadherin staining marks epithelial cell junctions.

**Supplementary Fig. 2:**
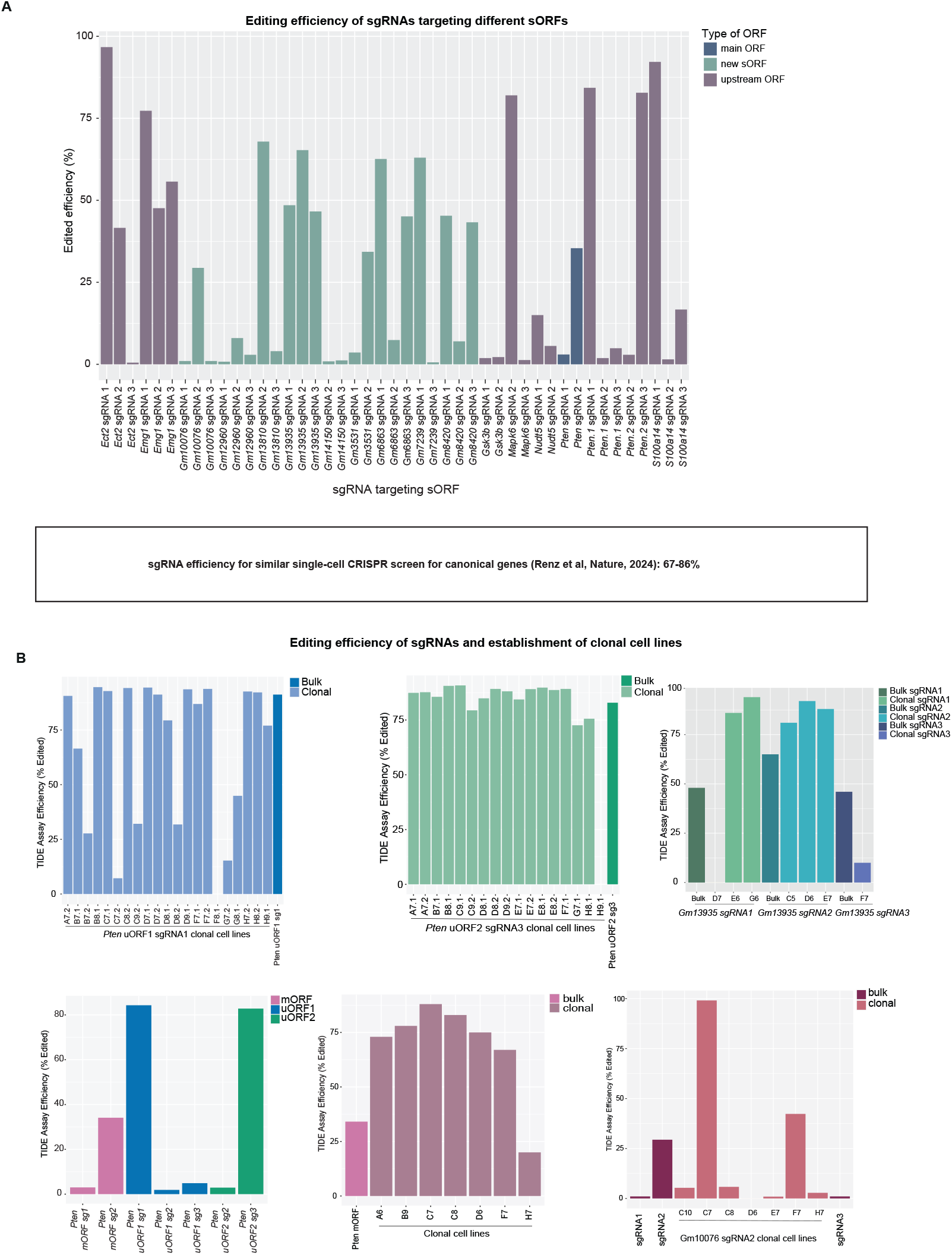
Gene editing efficiency of sgRNAs targeting microproteins. **(A)** sgRNA editing efficiency represented as the percentage of insertion and deletion (Indel) frequency. *Cas9*-positive keratinocytes were infected with the CROP-seq Puro vector containing single sgRNAs and selected for 7 days. Indel frequency was measured by the TIDE assay. Genomic locations ranging between 500 and 1200 base pairs were amplified from cells targeted with sgRNAs and the TIDE software was used to analyze the edits compared to non-targeted cells. Three sgRNAs per microprotein were tested for a total of 18 different microproteins. **(B)** Editing efficiency for clonal cells lines established for *Pten* uORF1, uORF2 and mORF as well as *Gm13935* and *Gm10076* clonal cell lines.

**Supplementary Fig. 3:**
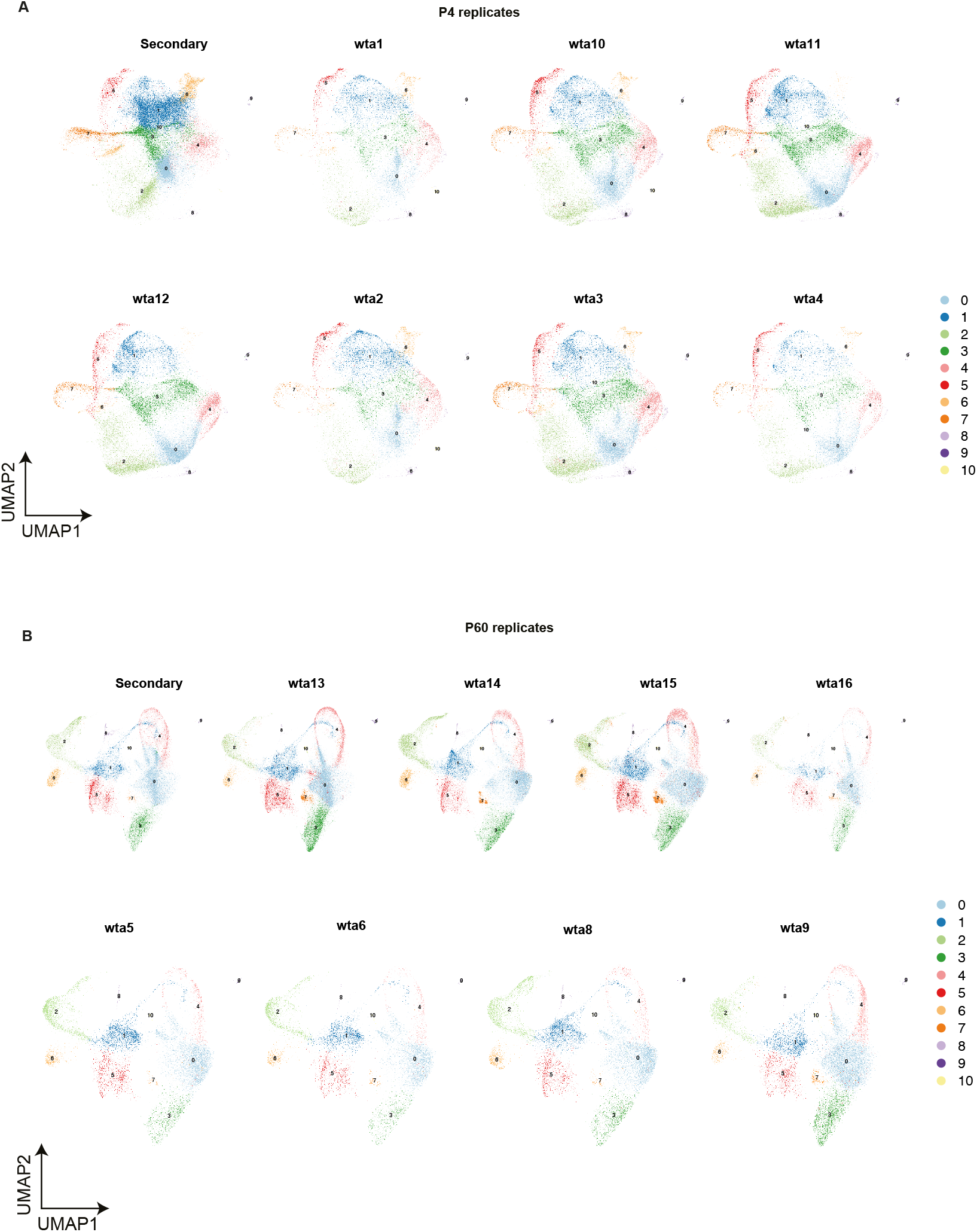
Single-cell CRISPR replicates from P4 and P60 epidermis exhibit consistent and uniform identification of cell types. **(A)** UMAPs for each P4 replicate for the primary and the secondary screen. All 11 cell types are detected in each of the replicates. **(B)** UMAPs for each P60 replicate for the primary and the secondary screen.

**Supplementary Fig. 4:**
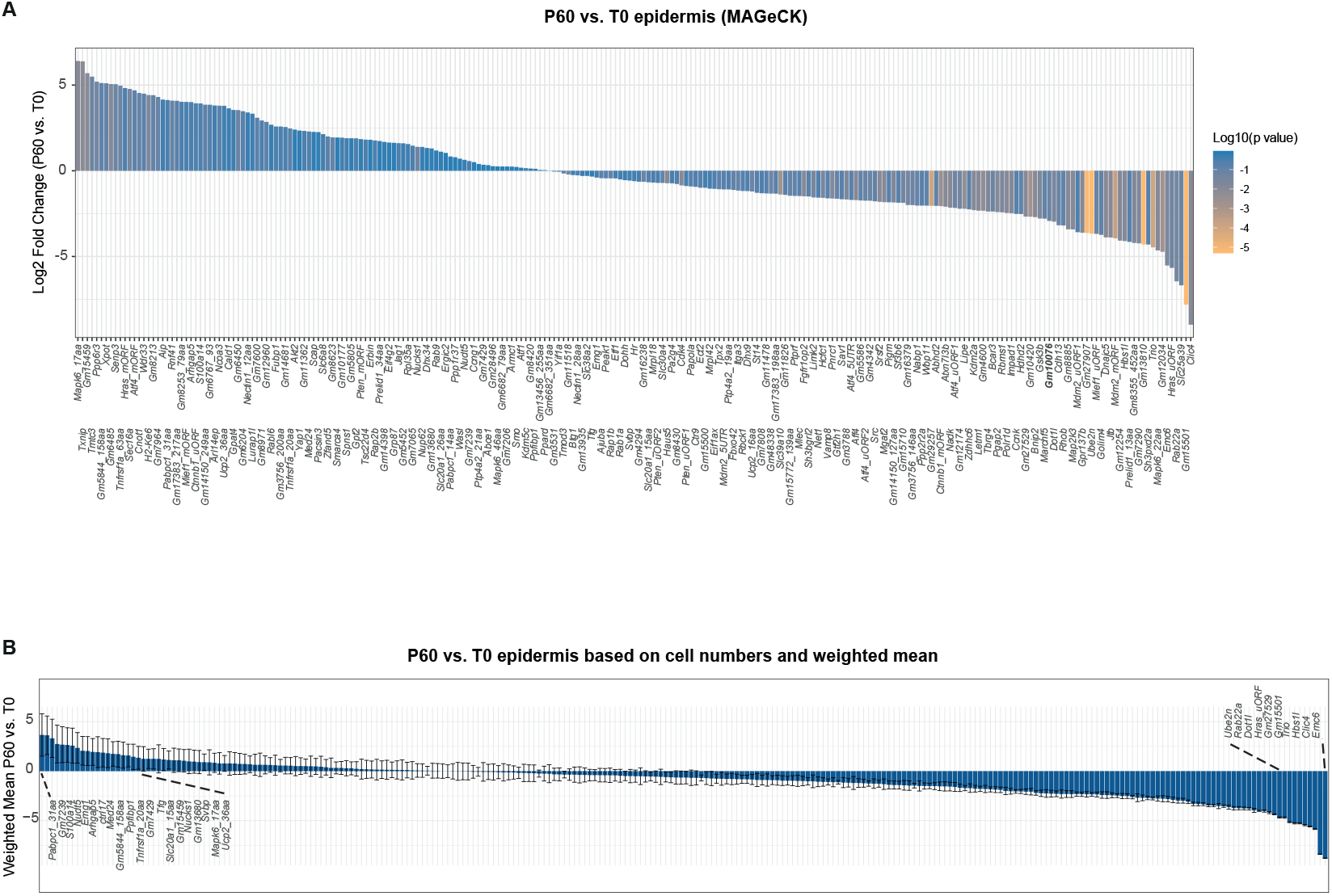
*In vivo* single-cell CRISPR screen in the P60 epidermis. **(A)** Waterfall plot showing the log2 fold change in sgRNA representation in the P60 epidermis compared to the library (T0), computed with MAGeCK and normalized to 50 nontargeting sgRNAs. Bars are colored by p value, highlighting the top enriched and depleted microprotein candidates. **(B)** Waterfall plot showing the log2 fold change in sgRNA representation based on cell numbers in the P60 epidermis compared to the library (T0) computed using weighted mean. The standard deviation is calculated as weighted approach, where the weight corresponds to the number of cells in each of the P60 replicate.

**Supplementary Fig. 5:**
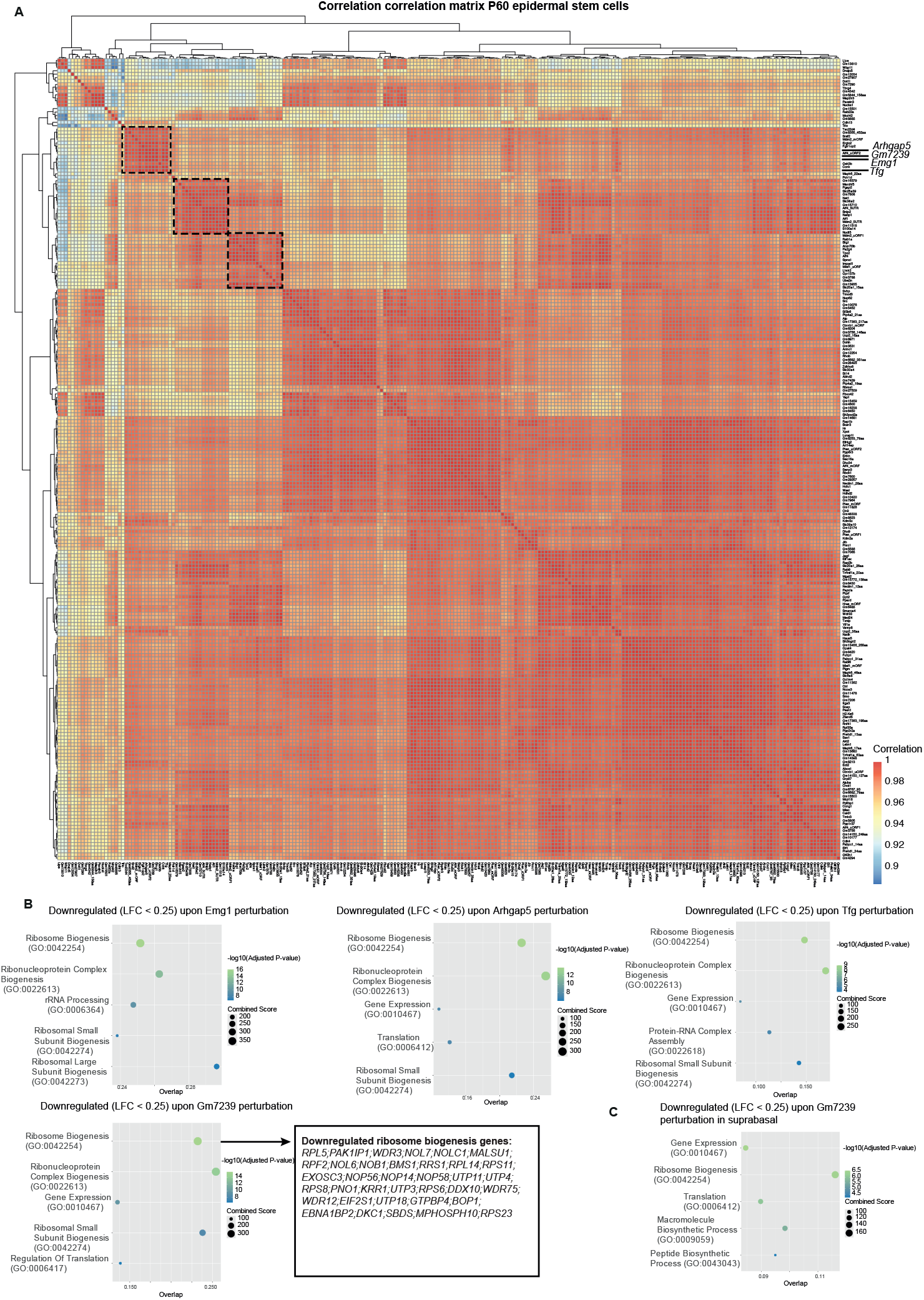
Clustering of perturbations in epidermal stem cells. **(A)** P60 perturbation-perturbation matrix based on gene expression in epidermal stem cells shows clustering of perturbations that lead to strong transcriptional changes (Fig. 3A, cohort of 14 perturbations), including *Emg1, Gm7239, Tfg* and *Arhgap5*. Perturbation-perturbation matrix of gene expression in P60 epidermal stem cells reveals clustering of perturbations that induce strong transcriptional changes (Fig. 3A, cohort of 14 perturbations), including *Emg1, Gm7239, Tfg* and *Arhgap5*. **(B)** Gene Ontology (GO) enrichment analysis of downregulated genes (p adjusted < 0.05; LFC < −0.25) in P60 epidermal stem cells highlights ribosome biogenesis affected by these perturbations. **(C)** GO enrichment analysis of downregulated genes in P60 suprabasal cells following *Gm7239* perturbation.

**Supplementary Fig. 6:**
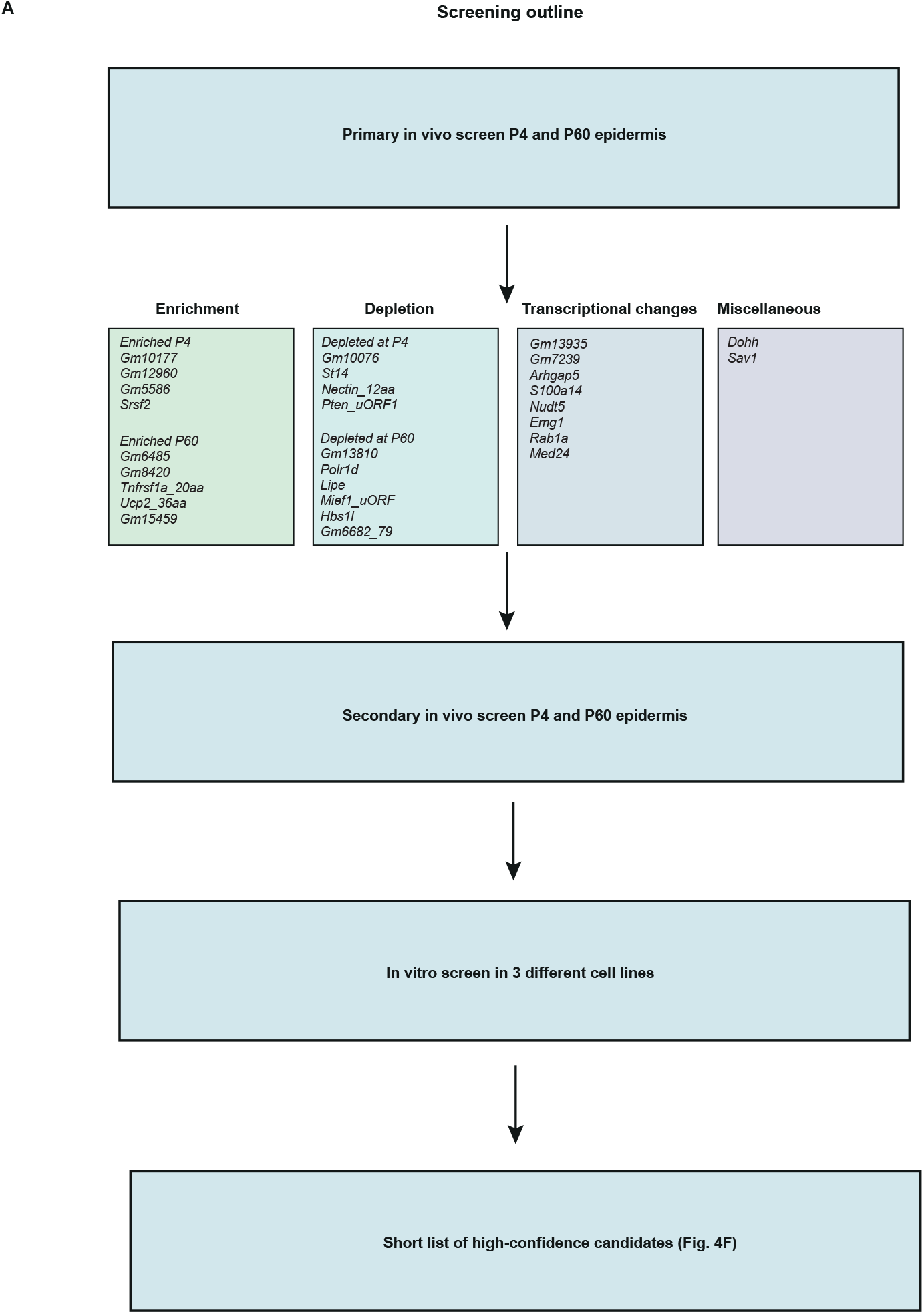
Selection of microprotein candidates for the validation screen. **(A)** Diagram outlining the criteria for selecting microprotein candidates for the *in vivo* and *in vitro* validation screens.

**Supplementary Fig. 7:**
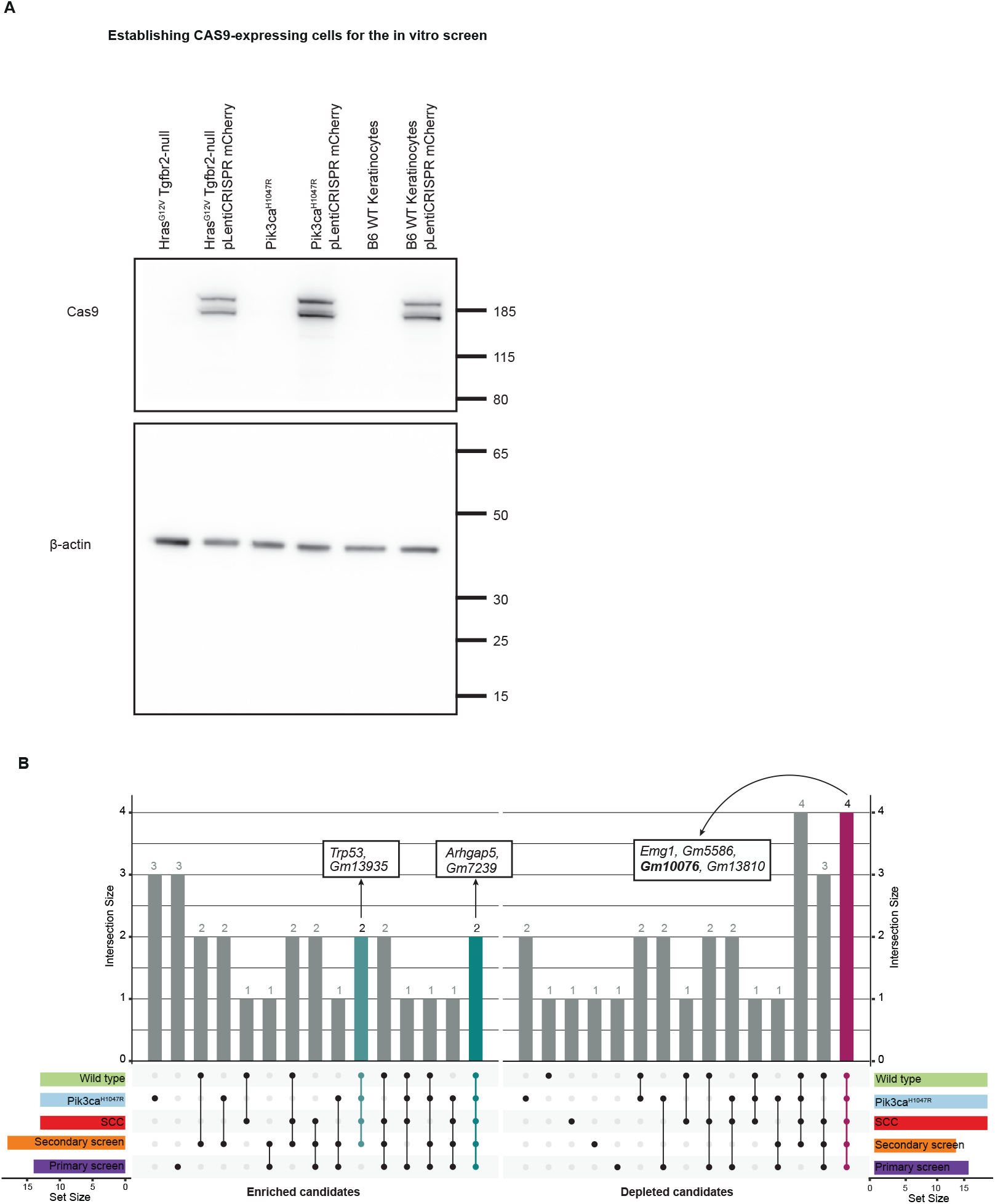
Establishing Cas9-expressing cells for the *in vitro* screen. **(A)** Western blot showing the 160 kDa Cas9 protein in keratinocytes infected with lentiviral pLentiCRISPR mCherry construct. *Hras*^*G12V*^; *Tgfbr2-null* (SCC cells), *Pik3ca*^*H1047*^ and wild-type keratinocytes are shown as controls next to the infected cells expressing Cas9. Lower panel shows the 45 kDa actin protein as a loading control for all cell types. **(B)** Upset plot summarizing the overlap of enriched and depleted microproteins across screens. *Arhgap5* and *Gm7239* are consistently enriched, while *Emg1, Gm5586, Gm10076* and *Gm13810* show consistent depletion across primary and validation screens.

**Supplementary Fig. 8:**
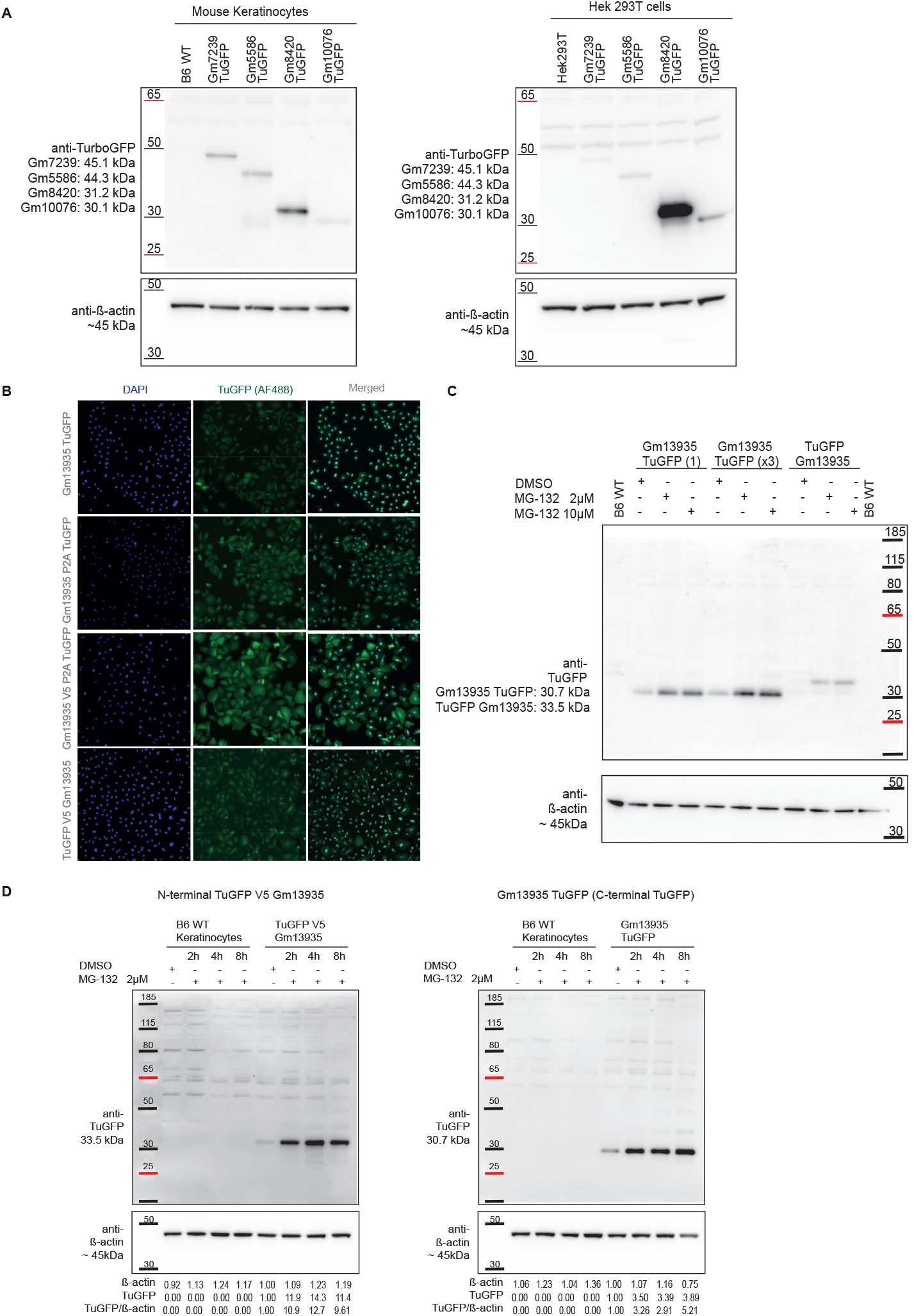
Characterization of candidate microproteins. **(A)** Western blot showing four microproteins tagged with Turbo GFP (TuGFP) in both wild-type keratinocytes and Hek293T cells. The following microproteins were tagged: *Gm7239::TuGFP* (45.1kDa), *Gm5586::TuGFP* (44.3kDa), *Gm8420::TuGFP* (31.2 kDa) and *Gm10076::TuGFP* (30.1 kDa). Lower panel shows the 45 kDa actin protein as loading control. **(B)** Immunofluorescence analyses of subcellular localization for the 31 amino acid *Gm13935* microprotein tagged with four different Turbo GFP constructs: *Gm13935::TuGFP, Gm13935::P2A::TuGFP, Gm13935::V5::P2A::TuGFP and TuGFP::V5::Gm13935*. **(C)** Western blot showing expression of C-terminally tagged *Gm13935::TuGFP* keratinocytes sorted sorted once or three times, alongside N-terminally tagged *TuGFP::V5::Gm13935* treated with the proteasome inhibitor MG132 (2 µM and 10 µM) for 2 hours. **(D)** Western blot showing a time-course analysis of C-terminally tagged Gm13935::TuGFP and N-terminally tagged TuGFP::V5::Gm13935 keratinocytes treated with proteosome inhibitor MG132 (2µM) for 2, 4 and 8 hours.

**Supplementary Fig. 9:**
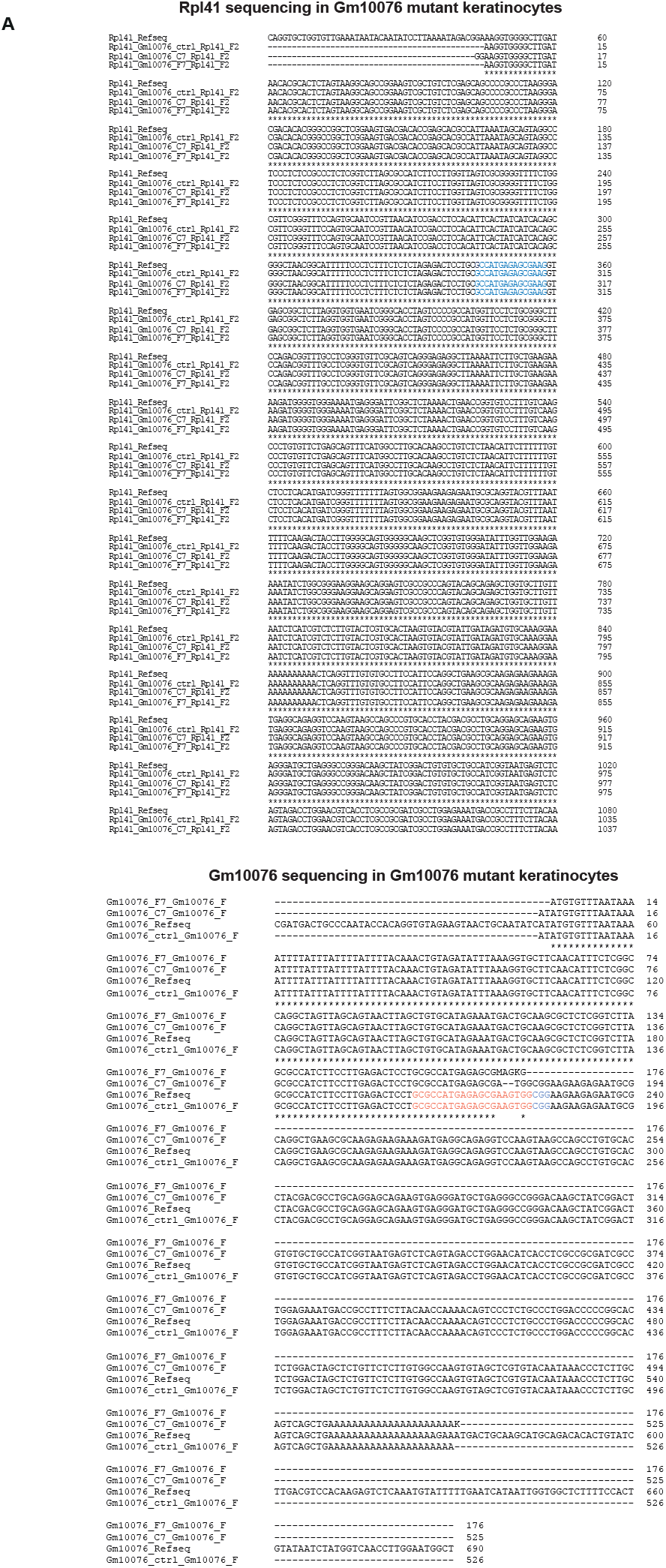
*Rpl41* and *Gm10076* locus sequencing in *Gm10076* mutant cells. **(A)** *Rpl41* is unedited in *Gm10076* mutant cells. Alignment of mutant *Gm10076* keratinocytes for the *Rpl41* gene on Chromosome 10. Two primer pairs flanking 816 nucleotides and 1154 nucleotides were designed to confirm that mutant *Gm10076* keratinocyte clonal lines (C7 and F7) remained unmodified compared to a control cell line. For *Rpl4*1, only 17 nucleotides of sgRNA2 aligned to the genomic region (highlighted in blue). Similarly, mutant *Gm10076* keratinocytes were tested for the *Gm10076* lncRNA genomic region on Chromosome 14, using primers amplifying 690 nucleotides flanking the predicted microprotein to verify genome editing in C7 and F7 clones. The sgRNA2 sequence is highlighted in orange, and the PAM site in blue. Clone C7 harbors a 2-nucleotide deletion, while clone F7 shows an interruption after the PAM site. Alignment results confirm that the *Gm10076* was targeted without altering the coding sequence of Rpl41.

**Supplementary Fig. 10:**
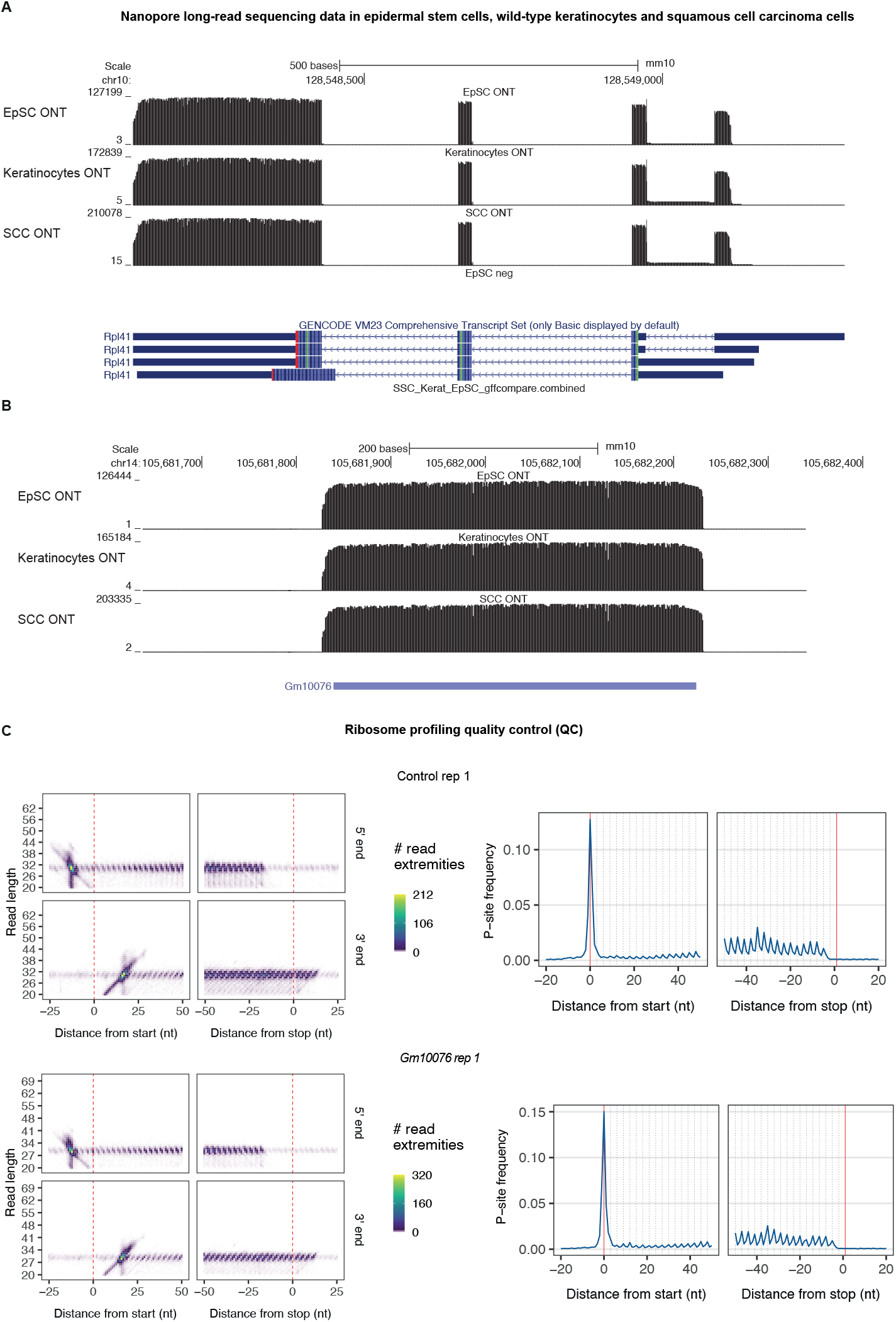
*Rpl41* and *Gm10076* are expressed at comparable levels in long-read sequencing data. **(A)** Nanopore long-read sequencing representation of the ribosomal gene *Rpl41*, displaying transcript annotation in epidermal stem cells, wild-type keratinocytes and squamous cell carcinoma cells. **(B)** Nanopore long-read sequencing representation of the lncRNA *Gm10076*, depicting transcript annotation across epidermal stem cells, wild-type keratinocytes and squamous cell carcinoma cells. Data retrieved from our previously published data set (https://genome-euro.ucsc.edu/s/umeshghosh/cage_ont) (*49*). **(C)** Quality control of ribosome profiling data, displaying metagene distribution of read length and P-site offset plots for control and *Gm10076 C7 KO* replicate 1.

**Supplementary Fig. 11:**
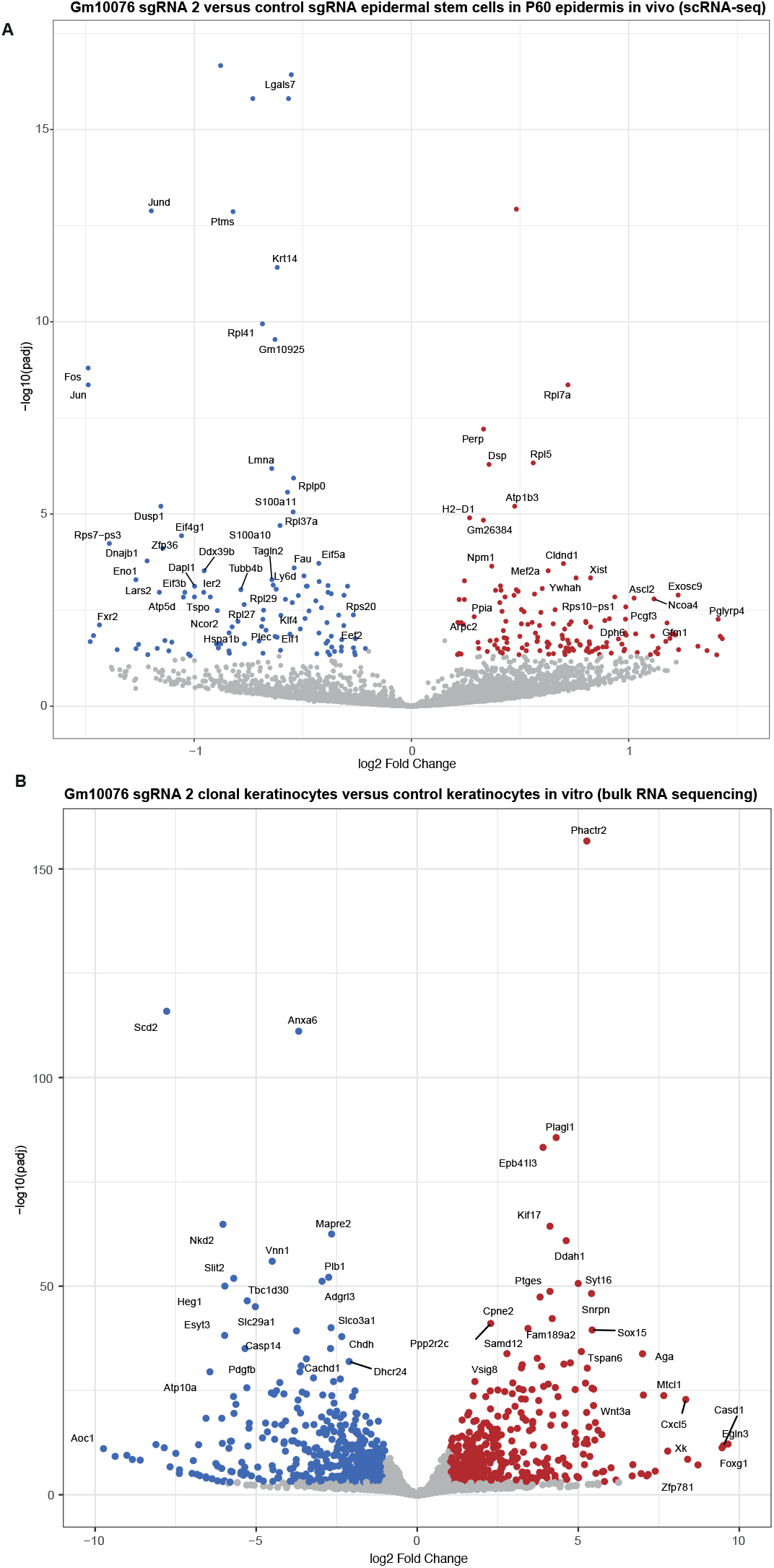
Transcriptional changes in *Gm10076* perturbed P60 epidermal stem cells and *Gm10076* mutant cells. **(A)** Volcano plot showing transcriptional changes in epidermal stem cells perturbed with *Gm10076 sgRNA2* compared to control sgRNA cells from the in vivo CRISPR screen (RNA-seq p adjusted < 0.05 and |LFC| > 0.25). **(B)** Volcano plot of transcriptional changes *Gm10076 C7* mutant keratinocytes compared to control keratinocytes (RNA-seq p adjusted < 0.05 and |LFC| > 0.25).

**Supplementary Fig. 12:**
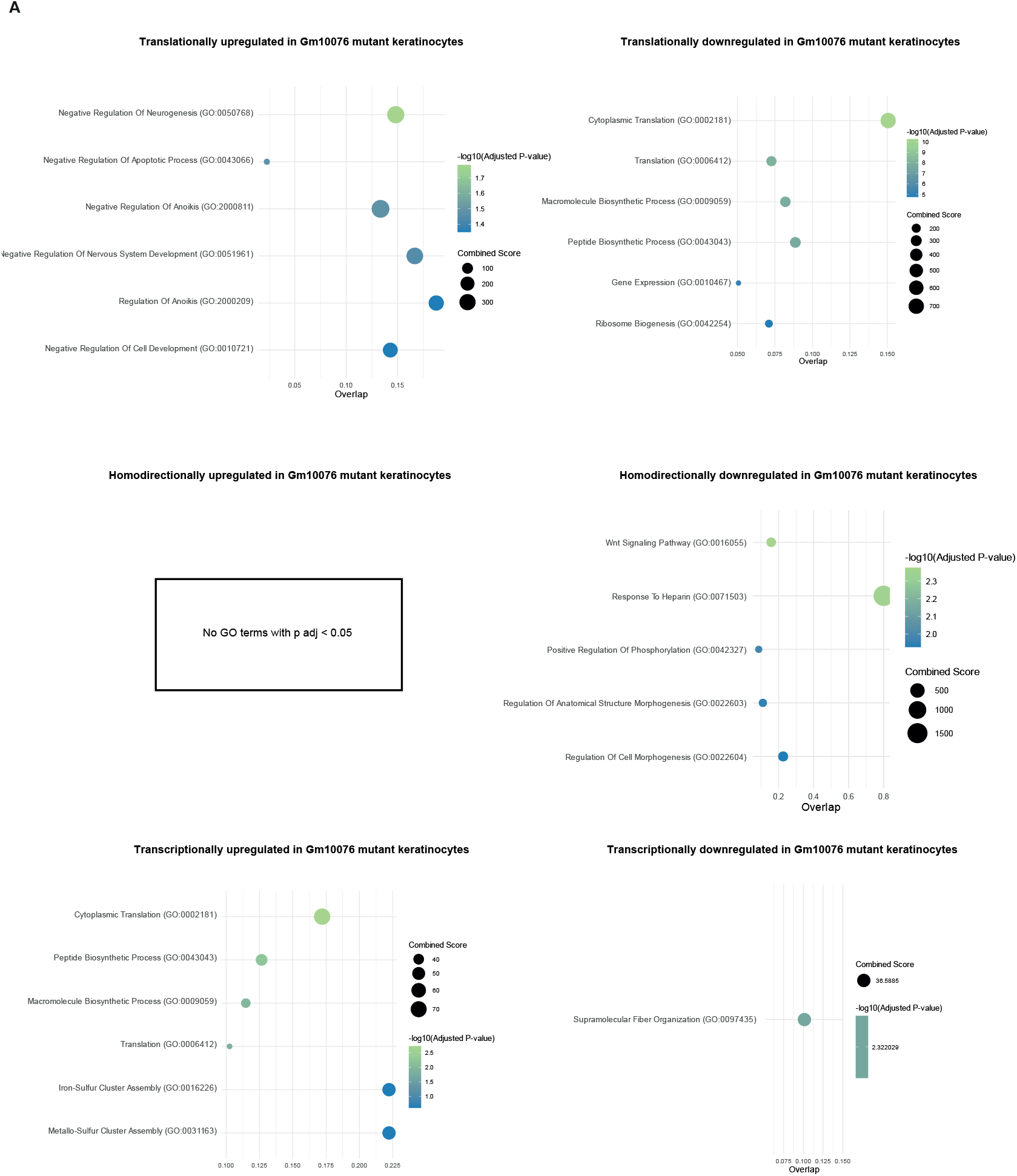
Gene ontology (GO) enrichment analysis upon *Gm10076* perturbation. **(A)** GO enrichment analysis for translationally upregulated and downregulated genes in *Gm10076* mutant keratinocytes. **(B)** GO enrichment analysis for homodirectionally upregulated and downregulated genes in *Gm10076* mutant keratinocytes. **(C)** GO enrichment analysis for transcriptionally upregulated and downregulated genes in *Gm10076* mutant keratinocytes.

**Supplementary Fig. 13:**
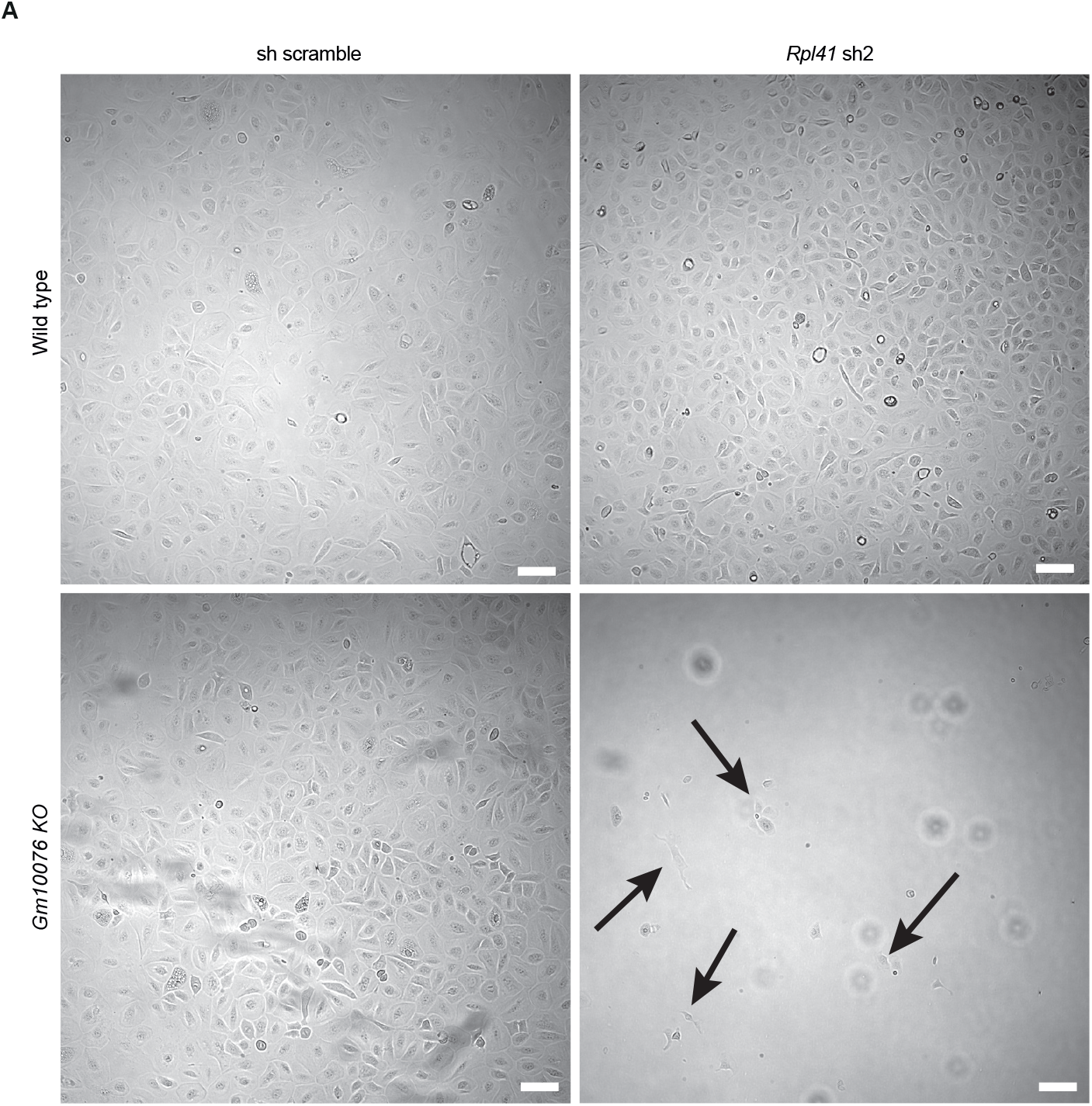
Knockdown of *Rpl41* in *Gm10076* mutant keratinocytes. **(A)** Representative images of *Rpl41 shRNA2; Gm10076 KO C7* two weeks post-infection with the lentiviral *Rpl41 shRNA2* construct. Arrows mark the few surviving *Rpl41 shRNA2; Gm10076 KO C7* keratinocytes. Scale bars, 100 μm.

**Supplementary Fig. 14:**
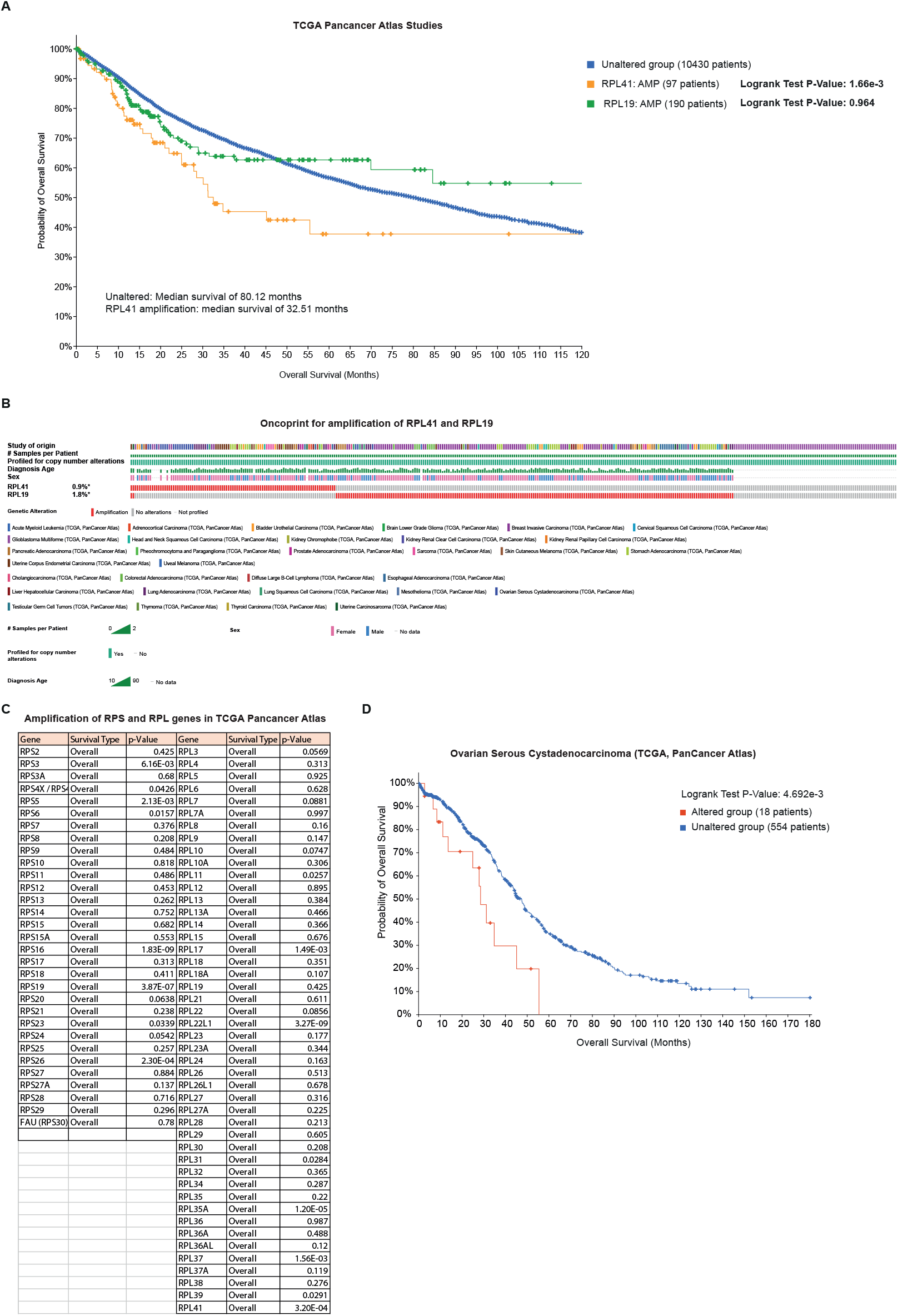
*RPL41* amplification is associated with shorter overall survival in cancer patients. **(A)** Analysis of TCGA PanCancer Atlas studies shows a significant correlation between RPL41 amplification and shorter overall survival. P values represent log-rank test results from cBioPortal. **(B)** Oncoprint visualization and detailed patient information for RPL41 and RPL19 amplifications in TCGA Pan-Cancer Atlas studies. **(C)** Analysis of ribosomal protein amplifications in TCGA PanCancer Atlas studies and their association with overall survival. P values represent log-rank test results from cBioPortal. **(D)** RPL41 amplification is associated with shorter overall survival in ovarian serous cystadenocarcinoma (TCGA). The p value represents the log-rank test value.

## References

1. J. M. Mudge, J. Ruiz-Orera, J. R. Prensner, M. A. Brunet, F. Calvet, I. Jungreis, J. M. Gonzalez, M. Magrane, T. F. Martinez, J. F. Schulz, Y. T. Yang, M.M. Albá, J. L. Aspden, P. V. Baranov, A. A. Bazzini, E. Bruford, M. J. Martin, L. Calviello, A.-R. Carvunis, J. Chen, J. P. Couso, E. W. Deutsch P. Flicek, A. Frankish, M. Gerstein, N. Hubner, N. T. Ingolia, M. Kellis, G. Menschaert, R. L. Moritz, U. Ohler, X. Roucou, A. Saghatelian, J. S. Weissman, S. van Heesch, Standardized annotation of translated open reading frames. Nat Biotechnol 40, 994–999 (2022).

2. E. W. Deutsch, L. W. Kok, J. M. Mudge, J. Ruiz-Orera, I. Fierro-Monti, Z. Sun, J. G. Abelin, M. M. Alba, J. L. Aspden, A. A. Bazzini, E. A. Bruford, M. A. Brunet, L. Calviello, S. A. Carr, A.-R. Carvunis, S. Chothani, J. Clauwaert, K. Dean, P. Faridi, A. Frankish, N. Hubner, N. T. Ingolia, M. Magrane, M. J. Martin, T. F. Martinez, G. Menschaert, U. Ohler, S. Orchard, O. Rackham, X. Roucou, S. A. Slavoff, E. Valen, A. Wacholder, J. S. Weissman, W. Wu, Z. Xie, J. Choudhary, M. Bassani-Sternberg, J. A. Vizcaíno, N. Ternette, R. L. Moritz, J. R. Prensner, S. van Heesch, High-quality peptide evidence for annotating non-canonical open reading frames as human proteins. bioRxiv, 2024.09.09.612016 (2024).

3. F. Valdivia-Francia, A. Sendoel, No country for old methods: New tools for studying microproteins. iScience 27, 108972 (2024).

4. D. A. Hofman, J. R. Prensner, S. van Heesch, Microproteins in cancer: identification, biological functions, and clinical implications. Trends Genet, S0168-9525(24)00211–7 (2024).

5. B. W. Wright, Z. Yi, J. S. Weissman, J. Chen, The dark proteome: translation from noncanonical open reading frames. Trends Cell Biol 32, 243–258 (2022).

6. J. J. Mohsen, A. A. Martel, S. A. Slavoff, Microproteins-Discovery, structure, and function. Proteomics 23, e2100211 (2023).

7. A. Saghatelian, J. P. Couso, Discovery and characterization of smORFencoded bioactive polypeptides. Nat Chem Biol 11, 909–916 (2015).

8. T. F. Martinez, Q. Chu, C. Donaldson, D. Tan, M. N. Shokhirev, A. Saghatelian, Accurate annotation of human protein-coding small open reading frames. Nat Chem Biol 16, 458–468 (2020).

9. C. Erady, A. Boxall, S. Puntambekar, N. Suhas Jagannathan, R. Chauhan, D. Chong, N. Meena, A. Kulkarni, B. Kasabe, K. Prathivadi Bhayankaram, Y. Umrania, A. Andreani, J. Nel, M. T. Wayland, C. Pina, K. S. Lilley, S. Prabakaran, Pan-cancer analysis of transcripts encoding novel open-reading frames (nORFs) and their potential biological functions. NPJ Genom Med 6, 4 (2021).

10. J. Chen, A.-D. Brunner, J. Z. Cogan, J. K. Nuñez, A. P. Fields, B. Adamson, D. N. Itzhak, J. Y. Li, M. Mann, M. D. Leonetti, J. S. Weissman, Pervasive functional translation of noncanonical human open reading frames. Science 367, 1140–1146 (2020).

11. D. A. Hofman, J. Ruiz-Orera, I. Yannuzzi, R. Murugesan, A. Brown, K. R. Clauser, A. L. Condurat, J. T. van Dinter, S. A. G. Engels, A. Goodale, J. van der Lugt, T. Abid, L. Wang, K. N. Zhou, J. Vogelzang, K. L. Ligon, T. N. Phoenix, J. A. Roth, D. E. Root, N. Hubner, T. R. Golub, P. Bandopadhayay, S. van Heesch, J. R. Prensner, Translation of non-canonical open reading frames as a cancer cell survival mechanism in childhood medulloblastoma. Mol Cell 84, 261–276.e18 (2024).

12. J. R. Prensner, O. M. Enache, V. Luria, K. Krug, K. R. Clauser, J. M. Dempster, A. Karger, L. Wang, K. Stumbraite, V. M. Wang, G. Botta, N. J. Lyons, A. Goodale, Z. Kalani, B. Fritchman, A. Brown, D. Alan, T. Green, X. Yang, J. D. Jaffe, J. A. Roth, F. Piccioni, M. W. Kirschner, Z. Ji, D. E. Root, T. R. Golub, Noncanonical open reading frames encode functional proteins essential for cancer cell survival. Nat Biotechnol 39, 697–704 (2021).

13. C. Zheng, Y. Wei, P. Zhang, K. Lin, D. He, H. Teng, G. Manyam, Z. Zhang, W. Liu, H. R. L. Lee, X. Tang, W. He, N. Islam, A. Jain, Y. Chiu, S. Cao, Y. Diao, S. Meyer-Gauen, M. Höök, A. Malovannaya, W. Li, M. Hu, W. Wang, H. Xu, S. Kopetz, Y. Chen, CRISPR-Cas9-based functional interrogation of unconventional translatome reveals human cancer dependency on cryptic non-canonical open reading frames. Nat Struct Mol Biol 30, 1878–1892 (2023).

14. D. Schlesinger, C. Dirks, C. Navarro, L. Lafranchi, A. Spinner, G. L. Raja, G. M.-S. Tong, J. Eirich, T. F. Martinez, S. J. Elsässer, A large-scale sORF screen identifies putative microproteins involved in cancer cell fitness. iScience 28 (2025).

15. P. F. Renz, U. Ghoshdastider, S. Baghai Sain, F. Valdivia-Francia, A. Khandekar, M. Ormiston, M. Bernasconi, C. Duré, J. A. Kretz, M. Lee, K. Hyams, M. Forny, M. Pohly, X. Ficht, S. J. Ellis, A. E. Moor, A. Sendoel, In vivo single-cell CRISPR uncovers distinct TNF programmes in tumour evolution. Nature 632, 419–428 (2024).

16. P. Datlinger, A. F. Rendeiro, C. Schmidl, T. Krausgruber, P. Traxler, J. Klughammer, L. C. Schuster, A. Kuchler, D. Alpar, C. Bock, Pooled CRISPR screening with single-cell transcriptome readout. Nat Methods 14, 297–301 (2017).

17. A. Sendoel, J. G. Dunn, E. H. Rodriguez, S. Naik, N. C. Gomez, B. Hurwitz, J. Levorse, B. D. Dill, D. Schramek, H. Molina, J. S. Weissman, E. Fuchs, Translation from unconventional 5′ start sites drives tumour initiation. Nature, doi: 10.1038/nature21036 (2017).

18. A. P. Fields, E. H. Rodriguez, M. Jovanovic, N. Stern-Ginossar, B. J. Haas, P. Mertins, R. Raychowdhury, N. Hacohen, S. A. Carr, N. T. Ingolia, A. Regev, J. S. Weissman, A Regression-Based Analysis of Ribosome-Profiling Data Reveals a Conserved Complexity to Mammalian Translation. Molecular Cell, doi: 10.1016/j.molcel.2015.11.013 (2015).

19. M. Fresno, A. Jiménez, D. Vázquez, Inhibition of Translation in Eukaryotic Systems by Harringtonine. European Journal of Biochemistry, doi: 10.1111/j.1432-1033.1977.tb11256.x (1977).

20. S. Beronja, E. Fuchs, RNAi-mediated gene function analysis in skin. Methods in Molecular Biology, doi: 10.1007/978-1-62703-227-8_23 (2013).

21. S. Beronja, P. Janki, E. Heller, W. H. Lien, B. E. Keyes, N. Oshimori, E. Fuchs, RNAi screens in mice identify physiological regulators of oncogenic growth. Nature, doi: 10.1038/nature12464 (2013).

22. M. S. Lawrence, C. Sougnez, L. Lichtenstein, K. Cibulskis, E. Lander, S. B. Gabriel, G. Getz, A. Ally, M. Balasundaram, I. Birol, R. Bowlby, D. Brooks, Y. S. N. Butterfield, R. Carlsen, D. Cheng, A. Chu, N. Dhalla, R. Guin, R. A. Holt, S. J. M. Jones, D. Lee, H. I. Li, M. A. Marra, M. Mayo, R. A. Moore, A. J. Mungall, A. Gordon Robertson, J. E. Schein, P. Sipahimalani, A. Tam, N. Thiessen, T. Wong, A. Protopopov, N. Santoso, S. Lee, M. Parfenov, J. Zhang, H. S. Mahadeshwar, J. Tang, X. Ren, S. Seth, P. Haseley, D. Zeng, L. Yang, A. W. Xu, X. Song, A. Pantazi, C. A. Bristow, A. Hadjipanayis, J. Seidman, L. Chin, P. J. Park, R. Kucherlapati, R. Akbani, T. Casasent, W. Liu, Y. Lu, G. Mills, T. Motter, J. Weinstein, L. Diao, J. Wang, Y. Hong Fan, J. Liu, K. Wang, J. Todd Auman, S. Balu, T. Bodenheimer, E. Buda, D. Neil Hayes, K. A. Hoadley, A. P. Hoyle, S. R. Jefferys, C. D. Jones, P. K. Kimes, Y. Liu, J. S. Marron, S. Meng, P. A. Mieczkowski, L. E. Mose, J. S. Parker, C. M. Perou, J. F. Prins, J. Roach, Y. Shi, J. V. Simons, D. Singh, M. G. Soloway, D. Tan, U. Veluvolu, V. Walter, S. Waring, M. D. Wilkerson, J. Wu, N. Zhao, A. D. Cherniack, P. S. Hammerman, A. D. Tward, C. Sekhar Pedamallu, G. Saksena, J. Jung, A. I. Ojesina, S. L. Carter, T. I. Zack, S. E. Schumacher, R. Beroukhim, S. S. Freeman, M. Meyerson, J. Cho, L. Chin, G. Getz, M. S. Noble, D. DiCara, H. Zhang, D. I. Heiman, N. Gehlenborg, D. Voet, P. Lin, S. Frazer, P. Stojanov, Y. Liu, L. Zou, J. Kim, C. Sougnez, S. B. Gabriel, M. S. Lawrence, D. Muzny, H. Doddapaneni, C. Kovar, J. Reid, D. Morton, Y. Han, W. Hale, H. Chao, K. Chang, J. A. Drummond, R. A. Gibbs, N. Kakkar, D. Wheeler, L. Xi, G. Ciriello, M. Ladanyi, W. Lee, R. Ramirez, C. Sander, R. Shen, R. Sinha, N. Weinhold, B. S. Taylor, B. Arman Aksoy, G. Dresdner, J. Gao, B. Gross, A. Jacobsen, B. Reva, N. Schultz, S. Onur Sumer, Y. Sun, T. A. Chan, L. G. Morris, J. Stuart, S. Benz, S. Ng, C. Benz, C. Yau, S. B. Baylin, L. Cope, L. Danilova, J. G. Herman, M. Bootwalla, D. T. Maglinte, P. W. Laird, T. Triche, D. J. Weisenberger, D. J. Van Den Berg, N. Agrawal, J. Bishop, P. C. Boutros, J. P. Bruce, L. Averett Byers, J. Califano, T. E. Carey, Z. Chen, H. Cheng, S. I. Chiosea, E. Cohen, B. Diergaarde, A. Marie Egloff, A. K. El-Naggar, R. L. Ferris, M. J. Frederick, J. R. Grandis, Y. Guo, R. I. Haddad, P. S. Hammerman, T. Harris, D. Neil Hayes, A. B. Y. Hui, J. Jack Lee, S. M. Lippman, F.-F. Liu, J. B. McHugh, J. Myers, P. Kwok Shing Ng, B. Perez-Ordonez, C. R. Pickering, M. Prystowsky, M. Romkes, A. D. Saleh, M. A. Sartor, R. Seethala, T. Y. Seiwert, H. Si, A. D. Tward, C. Van Waes, D. M. Waggott, M. Wiznerowicz, W. G. Yarbrough, J. Zhang, Z. Zuo, K. Burnett, D. Crain, J. Gardner, K. Lau, D. Mallery, S. Morris, J. Paulauskis, R. Penny, C. Shelton, T. Shelton, M. Sherman, P. Yena, A. D. Black, J. Bowen, J. Frick, J. M. Gastier-Foster, H. A. Harper, K. Leraas, T. M. Lichtenberg, N. C. Ramirez, L. Wise, E. Zmuda, J. Baboud, M. A. Jensen, A. B. Kahn, T. D. Pihl, D. A. Pot, D. Srinivasan, J. S. Walton, Y. Wan, R. A. Burton, T. Davidsen, J. A. Demchok, G. Eley, M. L. Ferguson, K. R. Mills Shaw, B. A. Ozenberger, M. Sheth, H. J. Sofia, R. Tarnuzzer, Z. Wang, L. Yang, J. Claude Zenklusen, C. Saller, K. Tarvin, C. Chen, R. Bollag, P. Weinberger, W. Golusiński, P. Golusiński, M. Ibbs, K. Korski, A. Mackiewicz, W. Suchorska, B. Szybiak, M. Wiznerowicz, K. Burnett, The Cancer Genome Atlas Network, Genome sequencing centre: Broad Institute, Genome characterization and data analysis centres: BC Cancer Agency, Harvard Medical School/Brigham & Women’s Hospital/MD Anderson Cancer Center, The University of Texas MD Anderson Cancer Center, University of Kentucky, University of North Carolina at Chapel Hill, Broad Institute, Baylor College of Medicine, Memorial Sloan-Kettering Cancer Center, University of California Santa Cruz/Buck Institute, Johns Hopkins University/Sidney Kimmel Comprehensive Cancer Center, University of Southern California, Disease working group, Biospecimen core resource: International Genomics Consortium, Nationwide Children’s Hospital, Data coordinating centre, Project office, Tissue source sites: Analytical Biological Services, Fred Hutchinson Cancer Research Center, Georgia Regents University, Greater Poland Cancer Centre, International Genomics Consortium, Comprehensive genomic characterization of head and neck squamous cell carcinomas. Nature 517, 576–582 (2015).

23. E. Cerami, J. Gao, U. Dogrusoz, B. E. Gross, S. O. Sumer, B. A. Aksoy, A. Jacobsen, C. J. Byrne, M. L. Heuer, E. Larsson, Y. Antipin, B. Reva, A. P. Goldberg, C. Sander, N. Schultz, The cBio Cancer Genomics Portal: An open platform for exploring multidimensional cancer genomics data. Cancer Discovery, doi: 10.1158/2159-8290.CD-12-0095 (2012).

24. S. K. Loganathan, K. Schleicher, A. Malik, R. Quevedo, E. Langille, K. Teng, R. H. Oh, B. Rathod, R. Tsai, P. Samavarchi-Tehrani, T. J. Pugh, A.-C. Gingras, D. Schramek, Rare driver mutations in head and neck squamous cell carcinomas converge on NOTCH signaling. Science 367, 1264–1269 (2020).

25. Z. Ying, M. Sandoval, S. Beronja, Oncogenic activation of PI3K induces progenitor cell differentiation to suppress epidermal growth. Nat Cell Biol 20, 1256–1266 (2018).

26. T. Jacob, K. Annusver, P. Czarnewski, T. Dalessandri, C. Kalk, C. Levra Levron, N. Campamà Sanz M. E. Kastriti, M. L. Mikkola, M. Rendl, B. M. Lichtenberger, G. Donati, Å. K. Björklund, M. Kasper, Molecular and spatial landmarks of early mouse skin development. Dev Cell 58, 2140–2162.e5 (2023).

27. S. Dekoninck, E. Hannezo, A. Sifrim, Y. A. Miroshnikova, M. Aragona, M. Malfait, S. Gargouri, C. de Neunheuser, C. Dubois, T. Voet, S. A. Wick-ström, B. D. Simons, C. Blanpain, Defining the Design Principles of Skin Epidermis Postnatal Growth. Cell 181, 604–620.e22 (2020).

28. I. Hollerer, J. C. Barker, V. Jorgensen, A. Tresenrider, C. Dugast-Dar-zacq, L. Y. Chan, X. Darzacq, R. Tjian, E. Ünal, G. A. Brar, Evidence for an Integrated Gene Repression Mechanism Based on mRNA Isoform Toggling in Human Cells. G3 (Bethesda) 9, 1045–1053 (2019).

29. H. Liang, S. He, J. Yang, X. Jia, P. Wang, X. Chen, Z. Zhang, X. Zou, M. A. McNutt, W. H. Shen, Y. Yin, PTENα, a PTEN Isoform Translated through Alternative Initiation, Regulates Mitochondrial Function and Energy Metabolism. Cell Metabolism 19, 836–848 (2014).

30. S.-M. Shen, C. Zhang, M.-K. Ge, S.-S. Dong, L. Xia, P. He, N. Zhang, Y. Ji, S. Yang, Y. Yu, J.-K. Zheng, J.-X. Yu, Q. Xia, G.-Q. Chen, PTENα and PTENβ promote carcinogenesis through WDR5 and H3K4 trimethylation. Nat Cell Biol 21, 1436–1448 (2019).

31. I. Tzani, I. P. Ivanov, D. E. Andreev, R. I. Dmitriev, K. A. Dean, P. V. Baranov, J. F. Atkins, G. Loughran, Systematic analysis of the PTEN 5’ leader identifies a major AUU initiated proteoform. Open Biol 6, 150203 (2016).

32. Z. Na, X. Dai, S.-J. Zheng, C. J. Bryant, K. H. Loh, H. Su, Y. Luo, A. F. Buhagiar, X. Cao, S. J. Baserga, S. Chen, S. A. Slavoff, Mapping subcellular localizations of unannotated microproteins and alternative proteins with MicroID. Mol Cell 82, 2900–2911.e7 (2022).

33. A. P. Wiita, E. Ziv, P. J. Wiita, A. Urisman, O. Julien, A. L. Burlingame, J. S. Weissman, J. A. Wells, Global cellular response to chemotherapy-induced apoptosis. eLife 2, e01236 (2013).

34. E.-H. Choi, S.-J. Park, TXNIP: A key protein in the cellular stress response pathway and a potential therapeutic target. Exp Mol Med 55, 1348–1356 (2023).

35. J. Deng, T. Pan, Z. Liu, C. McCarthy, J. M. Vicencio, L. Cao, G. Alfano, A. A. Suwaidan, M. Yin, R. Beatson, T. Ng, The role of TXNIP in cancer: a fine balance between redox, metabolic, and immunological tumor control. Br J Cancer 129, 1877–1892 (2023).

36. V. Justilien, K. C. Lewis, N. R. Murray, A. P. Fields, Oncogenic Ect2 signaling regulates rRNA synthesis in NSCLC. Small GTPases 10, 388–394 (2019).

37. P. Kalailingam, H. B. Tan, N. Jain, M. K. Sng, J. S. K. Chan, N. S. Tan, T. Thanabalu, Conditional knock out of N-WASP in keratinocytes causes skin barrier defects and atopic dermatitis-like inflammation. Sci Rep 7, 7311 (2017).

38. H. Li, S. Petersen, A. Garcia Mariscal, C. Brakebusch, Negative Regulation of p53-Induced Senescence by N-WASP Is Crucial for DMBA/TPA-Induced Skin Tumor Formation. Cancer Res 79, 2167–2181 (2019).

39. P. C. Liu, D. J. Thiele, Novel stress-responsive genes EMG1 and NOP14 encode conserved, interacting proteins required for 40S ribosome biogene-sis. Mol Biol Cell 12, 3644–3657 (2001).

40. J. Armistead, S. Khatkar, B. Meyer, B. L. Mark, N. Patel, G. Coghlan, R. E. Lamont, S. Liu, J. Wiechert, P. A. Cattini, P. Koetter, K. Wrogemann, C. R. Greenberg, K.-D. Entian, T. Zelinski, B. Triggs-Raine, Mutation of a gene essential for ribosome biogenesis, EMG1, causes Bowen-Conradi syn-drome. Am J Hum Genet 84, 728–739 (2009).

41. Q. Zhang, S. Wang, W. Li, E. Yau, H. Hui, P. K. Singh, V. Achuthan, M. A. Young Karris, A. N. Engelman, T. M. Rana, Genome-wide CRISPR/Cas9 transcriptional activation screen identifies a histone acetyltransferase inhibitor complex as a regulator of HIV-1 integration. Nucleic Acids Res 50, 6687–6701 (2022).

42. P. Wadsworth, TPX2. Curr Biol 25, R1156–1158 (2015).

43. G. Neumayer, C. Belzil, O. J. Gruss, M. D. Nguyen, TPX2: of spindle assembly, DNA damage response, and cancer. Cell Mol Life Sci 71, 3027–3047 (2014).

44. S. Klinge, F. Voigts-Hoffmann, M. Leibundgut, N. Ban, Atomic structures of the eukaryotic ribosome. Trends in Biochemical Sciences 37, 189–198 (2012).

45. A. Tsherniak, F. Vazquez, P. G. Montgomery, B. A. Weir, G. Kryukov, G. S. Cowley, S. Gill, W. F. Harrington, S. Pantel, J. M. Krill-Burger, R. M. Meyers, L. Ali, A. Goodale, Y. Lee, G. Jiang, J. Hsiao, W. F. J. Gerath, S. Howell, E. Merkel, M. Ghandi, L. A. Garraway, D. E. Root, T. R. Golub, J. S. Boehm, W. C. Hahn, Defining a Cancer Dependency Map. Cell 170, 564–576.e16 (2017).

46. J. Dresios, P. Panopoulos, K. Suzuki, D. Synetos, A Dispensable Yeast Ribosomal Protein Optimizes Peptidyltransferase Activity and Affects Translocation *. Journal of Biological Chemistry 278, 3314–3322 (2003).

47. T. C. Fleischer, C. M. Weaver, K. J. McAfee, J. L. Jennings, A. J. Link, Systematic identification and functional screens of uncharacterized proteins associated with eukaryotic ribosomal complexes. Genes Dev 20, 1294–1307 (2006).

48. N. Nagaraj, J. R. Wisniewski, T. Geiger, J. Cox, M. Kircher, J. Kelso, S. Pääbo, M. Mann, Deep proteome and transcriptome mapping of a human cancer cell line. Mol Syst Biol 7, 548 (2011).

49. R. Weber, U. Ghoshdastider, D. Spies, C. Duré, F. Valdivia-Francia, M. Forny, M. Ormiston, P. F. Renz, D. Taborsky, M. Yigit, M. Bernasconi, H. Yamahachi, A. Sendoel, Monitoring the 5′UTR landscape reveals isoform switches to drive translational efficiencies in cancer. Oncogene 42, 638–650 (2023).

50. I. Kisly, J. Remme, T. Tamm, Ribosomal protein eL24, involved in two intersubunit bridges, stimulates translation initiation and elongation. Nucleic Acids Res 47, 406–420 (2019).

51. M. I. Love, W. Huber, S. Anders, Moderated estimation of fold change and dispersion for RNA-seq data with DESeq2. Genome biology, doi: 10.1186/s13059-014-0550-8 (2014).

52. ‘Dark proteome’ survey reveals thousands of new human genes | Science | AAAS. https://www.science.org/content/article/dark-proteome-survey-reveals-thousands-new-human-genes.

53. Z. Zhang, P. Harrison, M. Gerstein, Identification and Analysis of Over 2000 Ribosomal Protein Pseudogenes in the Human Genome. Genome Res 12, 1466–1482 (2002).

54. P. Amaral, S. Carbonell-Sala, F. M. De La Vega, T. Faial, A. Frankish, T. Gingeras, R. Guigo, J. L. Harrow, A. G. Hatzigeorgiou, R. Johnson, T. D. Murphy, M. Pertea, K. D. Pruitt, S. Pujar, H. Takahashi, I. Ulitsky, A. Varabyou, C. A. Wells, M. Yandell, P. Carninci, S. L. Salzberg, The status of the human gene catalogue. Nature 622, 41–47 (2023).

55. E. W. Mills, R. Green, Ribosomopathies: There’s strength in numbers. Science 358, eaan2755 (2017).

56. M. Dodt, J. T. Roehr, R. Ahmed, C. Dieterich, FLEXBAR—Flexible Barcode and Adapter Processing for Next-Generation Sequencing Platforms. Biology (Basel) 1, 895–905 (2012).

57. Fast gapped-read alignment with Bowtie 2 | Nature Methods. https://www.nature.com/articles/nmeth.1923.

58. STAR: ultrafast universal RNA-seq aligner | Bioinformatics | Oxford Academic. https://academic.oup.com/bioinformatics/article/29/1/15/272537.

59. A. P. Fields, E. H. Rodriguez, M. Jovanovic, N. Stern-Ginossar, B. J. Haas, P. Mertins, R. Raychowdhury, N. Hacohen, S. A. Carr, N. T. Ingolia, A. Regev, J. S. Weissman, A Regression-Based Analysis of Ribosome-Profiling Data Reveals a Conserved Complexity to Mammalian Translation. Molecular Cell 60, 816–827 (2015).

60. P. Datlinger, A. F. Rendeiro, C. Schmidl, T. Krausgruber, P. Traxler, J. Klughammer, L. C. Schuster, A. Kuchler, D. Alpar, C. Bock, Pooled CRISPR screening with single-cell transcriptome readout. Nature Methods, doi: 10.1038/nmeth.4177 (2017).

61. S. Beronja, G. Livshits, S. Williams, E. Fuchs, Rapid functional dissection of genetic networks via tissue-specific transduction and RNAi in mouse embryos. Nat Med 16, 821–827 (2010).

62. C. Blanpain, W. E. Lowry, A. Geoghegan, L. Polak, E. Fuchs, Self-renewal, multipotency, and the existence of two cell populations within an epithelial stem cell niche. Cell 118, 635–648 (2004).

63. J. A. Nowak, E. Fuchs, Isolation and culture of epithelial stem cells. Methods Mol Biol 482, 215–232 (2009).

64. N. T. Ingolia, S. Ghaemmaghami, J. R. S. Newman, J. S. Weissman, Genome-wide analysis in vivo of translation with nucleotide resolution using ribosome profiling. Science, doi: 10.1126/science.1168978 (2009).

65. N. T. Ingolia, G. A. Brar, S. Rouskin, A. M. McGeachy, J. S. Weissman, The ribosome profiling strategy for monitoring translation in vivo by deep sequencing of ribosome-protected mRNA fragments. Nature Protocols, doi: 10.1038/nprot.2012.086 (2012).

66. N. J. MGlincy, N. T. Ingolia, Transcriptome-wide measurement of translation by ribosome profiling. Methods 126, 112–129 (2017).

67. Y. Hao, T. Stuart, M. H. Kowalski, S. Choudhary, P. Hoffman, A. Hartman, A. Srivastava, G. Molla, S. Madad, C. Fernandez-Granda, R. Satija, Dictionary learning for integrative, multimodal and scalable single-cell analysis. Nat Biotechnol 42, 293–304 (2024).

68. W. Li, H. Xu, T. Xiao, L. Cong, M. I. Love, F. Zhang, R. A. Irizarry, J. S. Liu, M. Brown, X. S. Liu, MAGeCK enables robust identification of essential genes from genome-scale CRISPR/Cas9 knockout screens. Genome Biology 15, 554 (2014).

69. C. Ahlmann-Eltze, W. Huber, glmGamPoi: fitting Gamma-Poisson generalized linear models on single cell count data. Bioinformatics 36, 5701–5702 (2021).

70. F. Lauria, T. Tebaldi, P. Bernabò, E. J. N. Groen, T. H. Gillingwater, G. Viero, riboWaltz: Optimization of ribosome P-site positioning in ribosome profiling data. PLOS Computational Biology 14, e1006169 (2018).

71. Z. Fang, X. Liu, G. Peltz, GSEApy: a comprehensive package for performing gene set enrichment analysis in Python. Bioinformatics 39, btac757 (2023).

